# Expression and subcellular localization of *USH1C*/harmonin in the human retina provide insights into pathomechanisms and therapy

**DOI:** 10.1101/2021.08.27.457962

**Authors:** Kerstin Nagel-Wolfrum, Benjamin R. Fadl, Mirjana M. Becker, Kirsten A. Wunderlich, Jessica Schäfer, Daniel Sturm, Jacques Fritze, Burcu Gür, Lew Kaplan, Tommaso Andreani, Tobias Goldmann, Matthew Brooks, Margaret R. Starostik, Anagha Lokhande, Melissa Apel, Karl R. Fath, Katarina Stingl, Susanne Kohl, Margaret M. DeAngelis, Ursula Schlötzer-Schrehardt, Ivana K. Kim, Leah A. Owen, Jan M. Vetter, Norbert Pfeiffer, Miguel A. Andrade-Navarro, Antje Grosche, Anand Swaroop, Uwe Wolfrum

**Affiliations:** Institute of Molecular Physiology, Johannes Gutenberg University Mainz, Germany; Institute of Developmental Biology and Neurobiology, Johannes Gutenberg University Mainz, Germany; Neurobiology, Neurodegeneration and Repair Laboratory, National Eye Institute, National Institutes of Health, Bethesda, USA; Department of Physiological Genomics, BioMedical Center, Ludwig-Maximilian University, Munich, Germany; Computational Biology and Data Mining, Institute of Organismic & Molecular Evolution Biology, Johannes Gutenberg University Mainz, Germany; Department of Ophthalmology, University Medical Centre Mainz, Germany; Queens College of CUNY, Kissena Blvd, Flushing, NY, USA; University Eye Hospital, Centre for Ophthalmology, University of Tubingen, Germany; Institute for Ophthalmic Research, Centre for Ophthalmology, University of Tubingen, Germany; Department of Ophthalmology and Ira G. Ross Eye Institute, Jacobs School of Medicine and Biomedical Sciences, University of Buffalo, USA; Department of Ophthalmology, Friedrich-Alexander-Universität, Erlangen-Nürnberg, Germany; Retina Service, Massachusetts Eye and Ear Infirmary, Harvard Medical School, Boston, USA; Department of Ophthalmology and Visual Sciences, University of Utah, Salt Lake City, USA

**Keywords:** Usher syndrome, retina dystrophy, ciliopathy, USH1C, harmonin, alternate splicing, splicing variants, photoreceptor cells, Müller glia cell, cell-cell adhesions, outer limiting membrane

## Abstract

Usher syndrome (USH) is the most common form of hereditary deafness-blindness in humans. USH is a complex genetic disorder, assigned to three clinical subtypes differing in onset, course, and severity, with USH1 being the most severe. Rodent USH1 models do not reflect the ocular phenotype observed in human patients to date; hence, little is known about the pathophysiology of USH1 in the human eye. One of the USH1 genes, *USH1C*, exhibits extensive alternative splicing and encodes numerous harmonin protein isoforms that function as scaffolds for organizing the USH interactome. RNA-seq analysis of human retinas uncovered harmonin_a1 as the most abundant transcript of *USH1C*. Bulk RNA-seq analysis and immunoblotting showed abundant expression of harmonin in Müller glia cells (MGCs) and retinal neurons. Furthermore, harmonin was localized in the terminal endfeet and apical microvilli of MGCs, presynaptic region (pedicle) of cones, and outer segments of rods as well as at adhesive junctions of MGCs and photoreceptors in the outer limiting membrane (OLM). Our data provide evidence for the interactions of harmonin with OLM molecules in photoreceptors (PRCs) and MGCs and rhodopsin in PRCs. Subcellular expression and colocalization of harmonin correlate with the clinical phenotype observed in USH1C patients. In addition, primary cilia defects in *USH1C* patient-derived fibroblasts could be reverted by the delivery of harmonin_a1 transcript isoform. Our data provide novel insights into PRC cell biology, *USH1C* pathophysiology, and for developing gene therapy treatment.

## Introduction

Usher syndrome (USH) is a clinically and genetically heterogeneous disease and the most frequent cause of deaf-blindness in humans affecting one in 6-10,000 individuals (1, 2). USH is considered a retinal ciliopathy because of defects in photoreceptor cilia and other primary cilia (3-5). Among the three clinical subtypes of USH (USH1, USH2, and USH3) USH1 is the most severe, characterized by profound congenital hearing impairment or deafness, vestibular dysfunction, and prepuberal onset of progressive retinal degeneration (6).

As of now, for USH1 six genes have been identified (6), which encode proteins of different protein families that collectively function in dynamic networks at distinct subcellular locations in the eye and the inner ear (7). The human *USH1C* gene (ENSG00000006611; OMIM 276904) is located on chromosome 11 (chr 11:17,493,895-17,544,416 on GRCh38.p10) and encodes the harmonin protein. The gene consists of 28 exons spanning over 50 kb of genomic region (Figure 1A), and 11 distinct transcripts (splice variants) have been annotated in humans, so far (Ensembl ENSG00000006611) (8).

**Figure 1.**
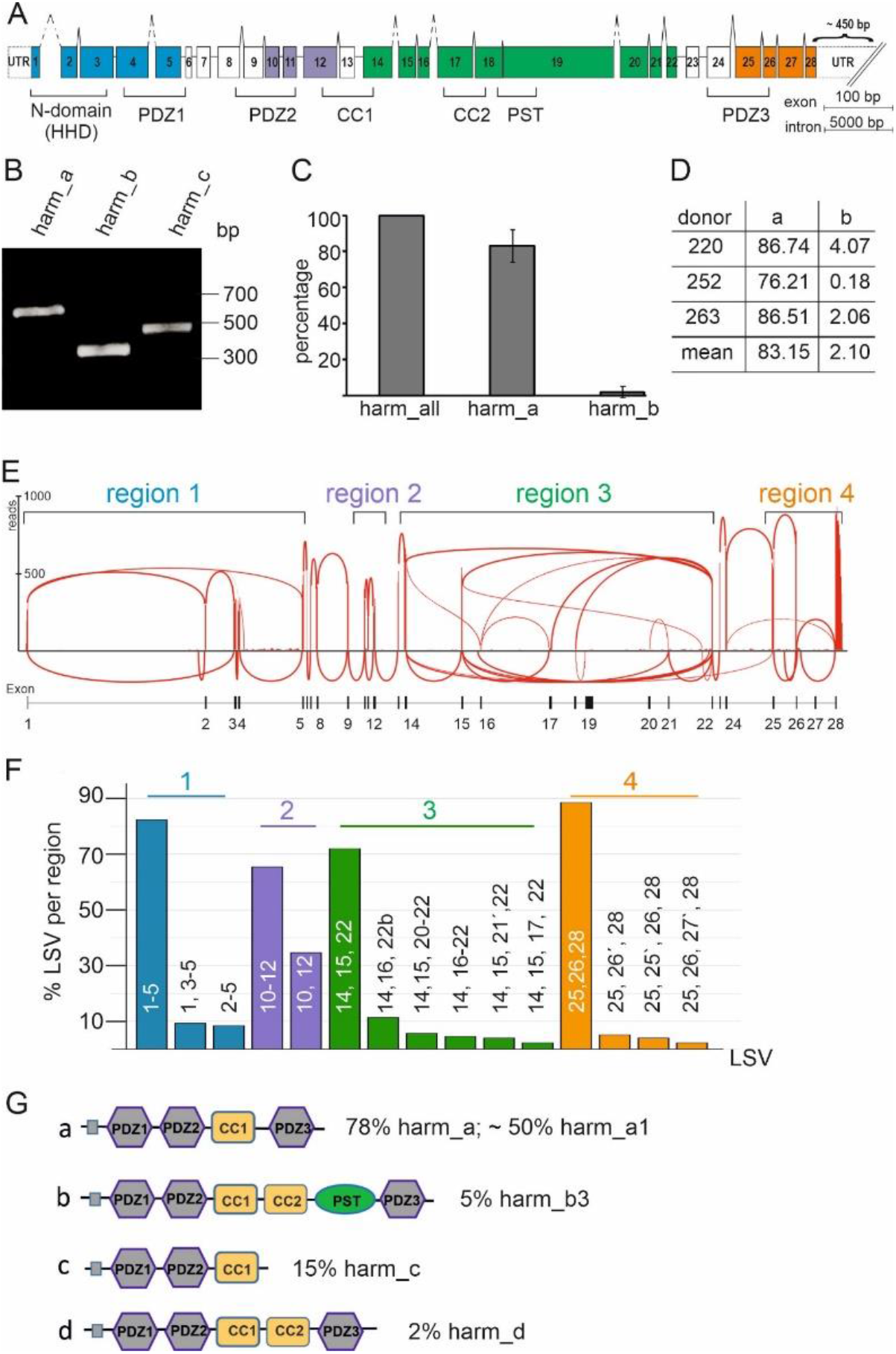
Expression of *USH1C*/harmonin a, b and c transcripts in the human retina. (**A**) Exon and intron structure of human *USH1C*. Exons are shown in boxes and denoted by the numbers from 1 to 28. The different domains of the harmonin protein, namely *N-*terminal or harmonin homology domain (HHD), PDZ (PSD-95, DLG, ZO-1) domains, PST (proline-serine-threonine rich) domain, CC (coiled-coil) domains, are marked by brackets below the gene structure. Colors indicate regions of local splice variant (LSV) analyses. (**B-D**) RT-PCR analyses of *USH1C*/harmonin transcripts using isoform specific primers in adult human retina. (**B**) RT-PCR analysis of *USH1C*/harmonin transcripts using isoform specific primers in human retina. Transcripts of *USH1C*/harmonin isoforms a, b and c were detected. Equal amounts of amplified cDNA were loaded. (**C**) Quantitative RT-PCR of *USH1C*/harmonin isoforms. Primers detect either all *USH1C*/harmonin transcripts (a, b and c) or are specific for *USH1C*/harmonin_a or *USH1C*/harmonin_b transcripts, respectively. *USH1C*/harmonin_a isoforms are most prominent, while *USH1C*/harmonin_b isoforms are rarely expressed. (**D**) Percentages of transcripts in three different donors are shown. *USH1C*/harmonin_a-transcripts were most abundant (83.15%), *USH1C*/harmonin_b-transcripts were barely detected (2.1%). **(E)** Representative Sashimi plot of RNA-seq analysis of human retina. Splicing events were grouped into four regions (region 1-4). **(F)** Quantification of local splice variants (LSV). LSVs of the same region are depicted in same color. (**G)** Predicted domain structures of the *USH1C*/harmonin transcripts. Percentage of transcripts is based on the copy number of LSVs in regions 1-4. *USH1C*/harmonin class a is the most abundant class with *USH1C*/ harmonin_a1 being the most expressed isoform.

*Ush1c* transcripts in murine retinae exhibit extensive alternative splicing (9) with at least nine different isoforms encoding three different classes (a, b and c), classified by their domain contents in the expressed harmonin protein (Figure 1) (8, 10) (www.ensembl.org). Characteristic sequences in all harmonin isoforms are 90-100 amino acid long PDZ domains, named for the three scaffold proteins originally recognized to contain these sequences: <u>P</u>SD-95, <u>D</u>LG, and <u>Z</u>O-1 (11). PDZs contain a hydrophobic pocket that mainly associates with PDZ-binding motifs (PBM) present mostly at the *C*-terminus of target proteins. A majority of the interacting partners of harmonin bind to as many as three PDZ domains, making it a potent scaffold protein with a central position in the USH interactome (8, 12). In addition to myosin VIIa (USH1B), cadherin 23 (USH1D), and SANS (USH1G), the USH2 proteins USH2A and VLGR1 (USH2C) bind to PDZ1 domain in harmonin, whereas cadherin 23 (USH1D) and protocadherin 15 (USH1F) bind to the PDZ2(8). SANS additionally interacts with PDZ3. Notably, all harmonin isoforms contain a CC (coiled-coil) domain and a conserved globular *N*-terminal domain also named as the harmonin homology domain (HHD) (13), which increases the affinity of proteins binding to PDZ1, such as SANS, VLGR1 or USH2A (14-16). Harmonin_b, the longest isoform, is characterized by a second CC domain followed by a PST (proline-serine-threonine rich) domain that facilitates actin filament bundling (17). In addition, harmonin can interact with non-USH proteins in specific cellular compartments; e.g., the subunits of voltage-gated Ca^2+^-channels at ribbon synapses (18, 19), and the GTPase regulator DOCK4 in stereocilia (20).

Alternative splicing of *USH1C* transcripts further increases the functional properties of the harmonin scaffold protein providing functional diversity to the harmonin’s interaction profile in relevant cells. *USH1C*/harmonin isoforms are expressed in a wide range of tissues, yet studies on expression and subcellular localization of harmonin in rodent retinae have not been definitive (10, 15, 21-23). Based on the depicted subcellular localization of harmonin isoforms in murine photoreceptor cells (PRCs), others and we have suggested differential scaffold functions of harmonin especially in the outer segment and at the synapses (8, 15, 21, 24, 25). A putative synaptic function has also been indicated in the murine retina (22), confirmed by zebrafish morpholino knock-downs (26) and recently by the USH1C pig model (27).

In the human retina, harmonin has been additionally found along with other USH proteins in the calyceal processes of cone and rod PRCs (23). These extensions of the PRC inner segment are suggested to stabilize the light-sensitive outer segment of the cell against mechanical forces (4, 28). Interestingly, the PRCs of most vertebrates, except for rodents, possess calyceal processes. Conclusively, absence of calyceal processes in mice is thought to be the reason why USH1 mouse models do not have a retinal phenotype comparable to human USH1 patients (23, 28).

Previous studies on the retina indicated that *USH1C/*harmonin is mainly expressed in PRCs and to a lesser degree in secondary retinal neurons (8, 21-23). However, harmonin was also shown to be expressed in Müller glia cells (MGCs) (26). More recent single cell (sc)RNA-seq studies of human retinal cells even indicated that the expression of *USH1C* is almost exclusively restricted to MGCs (29, 30). MGCs are the radial glial cells of the retina extending from the outer limiting membrane through all retinal layers to the extracellular inner limiting membrane that separates the neural retina from the vitreous body of the eye. Functionally, MGCs not only maintain the structural integrity of the retina but are also essential for retinal homeostasis and physiology (31, 32). However, nothing is known currently about the subcellular localization and function of harmonin in the MGCs.

Development of ocular gene augmentation therapies require a detailed understanding of the expression profile of *USH1C*/harmonin to delineate which isoform and into what retinal cell types this has to be introduced (33). In this study, we investigated the expression of *USH1C*/harmonin at both the transcript and the protein level in the human retina, by RNA-seq and Western blotting as well as correlative and *in situ* imaging techniques. We describe extensive splicing of *USH1C* and identify harmonin_a1 as the most prominent isoform in the human retina. We demonstrate *USH1C*/harmonin expression and subcellular localization in MGCs as well as in retinal rod and cone PRCs. In addition, we provide evidence for novel interacting proteins of harmonin in distinct subcellular compartments. Finally, we identified altered cilia length in fibroblasts from USH1C patients as an important phenotype and demonstrate that we can reverse the pathogenic phenotype by addition of harmonin_a1 to patient-derived cells.

## Results

### *USH1C* is frequently spliced in the human retina

To elaborate the transcriptomic profile of *USH1C* in the human retina, we first validated the expression of all three reported classes of *USH1C/*harmonin isoforms by RT-PCR in five healthy human donor retinae (Figure 1B-D, Supplemental Tables S1, S2, and S3). RT-PCR demonstrated amplicons specific for all three isoform classes: a, b, and c (Figure 1B). Subsequent qRT-PCRs of human donor retinas revealed the proportional presence of ∼83% USH1C/harmonin_a and ∼2% USH1C/harmonin_b splice variants of total *USH1C*/harmonin expression in human retina (Figure 1C,D). The remaining isoform classes (c and d, see below) are likely to account for approximately 15% of *USH1C* transcripts.

Next, we analyzed the local splicing events of *USH1C* in a human retina using RNA-seq. We generated an RNA-seq library of the human retina with approximately 90 million reads and 151bp paired-end read-length and analyzed aligned RNA-seq data using Sashimi plots (34) (Figure 1E). We observed extensive alternative splicing of *USH1C* in the human retina. For the analysis of the splicing events that are occurring on *USH1C*, we grouped the splicing events into four regions, namely exons 1 to 5 (region 1), exons 10 to 12 (region 2), exons 14 to 22 (region 3), and exons 25 to 28 (region 4), which were separated by canonically spliced exon-exon junctions (Figure 1E). For the quantification of the observed local splice variants (LSV), we utilized 50 additional published RNA-seq samples (35). These samples had a comparably low sequencing depth of ∼10 million reads per sample and rarely occurring LSVs which would appear as random splice events. Hence, we consolidated the 50 samples in 10 samples by iteratively random sampling reads from each of the 50 samples which eventually were pooled to avoid the probability that a splice variant observed was driven only by some of the samples. This procedure made it possible to obtain a total of 10 fused samples with homogeneous numbers of reads from the 50 original samples with low coverage, in order to increase coverage for the detection of potential splice variants. We then quantified each observed LSV in the merged samples. Only LSVs with a minimum of 20 raw counted reads in at least half of the merged samples were used for any further subsequent downstream analysis.

The LSV ex(1-5) is the most frequent LSV in region 1 of *USH1C* with ∼82% read coverage within region 1 (Figure 1F). A novel LSV ex(1,3-5), in which exon 2 is skipped (∼9% read coverage), leads to a frameshift and a premature stop codon in exon 3. LSV ex(2-5) is characterized by an alternative start and had an abundance of ∼8% among all LSVs in region 1.

In region 2 (exons 10-12), we observed two LSV which differ in the presence or absence of exon 11. Exon 11 was present in about two thirds of the reads. LSV ex(10,12) does not shift the open reading frame (ORF) and can be translated into a *USH1C/*harmonin isoform with a slightly shortened PDZ2 domain. Both LSVs are known and annotated transcripts for *USH1C/*harmonin_a1 (ENST00000318024) and a4 (ENST00000527020), respectively.

The alternative splicing in region 3 (exons 14-22) was very complex. At the protein level, this region encodes for the CC2 and PST domains, only present in *USH1C/*harmonin class_b isoforms. The most frequent LSV in region 3 was LSV ex(14,15,22) with ∼72% read coverage (Figure 1F). This LSV lacks the CC2 and PST domains and therefore encodes for *USH1C/*harmonin class_a or _c. LSV ex(14,16-22) was only covered with a very low number of reads in the RNA-seq samples. However, this LSV encodes for the CC2 and PST domains indicating a low expression of *USH1C/*harmonin isoform_b in the human retina. Thus, these RNA-seq data confirmed the qRT-PCR results (Figure 1B-D). We further found two potentially new exons (16’ and 21’) located between exons 16 and 17, and 21 and 22, respectively, and a putative alternative 5’ splice site of exon 22.

Detailed analysis in region 4 revealed three LSVs. LSV ex(25,26,28) is the most frequent splicing event with a read coverage of ∼90%. Two novel exons (exons 25’ and 26’) which have not been previously described for *USH1C* were also observed (Figure 1F). Exon 25’ was part of LSV ex(25,25’,26,28) and exon 26’ was part of LSV ex(25,26’,28), both with a read coverage of ∼5%.

To translate our computational analyses, we combined the most abundant LSVs of regions 1, 2 and 4 with each LSV found in region 3 and subsequently used these in the online tool SMART (http://SMART.embl-heidelberg.de) to predict the domain structure (Figure 1G). Most LSVs were assigned to the *USH1C*/harmonin classes a, b and c (8, 10). In addition, we found a putative novel class for harmonin, which we named *USH1C*/harmonin_d that is similar to harmonin_a, but additionally carries a CC2 domain (Figure 1G). Approximately ∼78% of all LSVs in region 3 were predicted to result in *USH1C/*harmonin_a isoforms, of which approximately two thirds are *USH1C/*harmonin_a1 and the remaining harmonin_a4 depending on the splicing of exon 11 (region 2). This quantification indicated that ∼50% of all harmonin expressed in the human retina is *USH1C/*harmonin_a1

### *USH1C* is highly expressed in both Müller glia cells and retinal neurons

To assign *USH1C* transcripts to specific cell types of the human retina, we performed bulk RNA-seq from sorted MGCs and retinal neurons (RNs) from human donor retinae (Supplemental Tables S1 and S2). As described above, we grouped the splicing events into four regions and set a cut-off of 5% for each LSV per region (Figure 2A). In all four regions *UHS1C*/harmonin transcripts were identified in RNs and MGCs (Supplemental Table S2). In region 1, LSV ex(1-5) was the most frequent LSV, for both RNs and MGCs. Interestingly, expression of LSV ex(1-5) was increased in MGCs compared to RNs. In addition to LSV ex(1,3-5) and LSV ex(2-5), which were both expressed in the two cell types, we identified LSV ex(1,5) in RNs only. In region 2, we observed the two LSVs (ex(10-12; ex(10,12)having no differences in the number of LSV between RNs and MGCs. In region 3, our bulk RNA-seq revealed 3 LSVs,. namely the most frequent transcript LSV ex(14,15,22), being slightly higher expressed in MGCs compared to RNs. LSV ex(14,15,20-22) and LSV ex(14,15,21,22), encoding for the *C-*terminal part of the PST domain were detected, too. However, in contrast to the RNA-seq of the total retina, no LSV ex(18,19) encoding the entire PST domain was detected, probably due to lower sequencing depth. In region 4, LSV ex(25,26,28) was the most abundant LSV and expressed in both RNs and MGCs.

**Figure 2.**
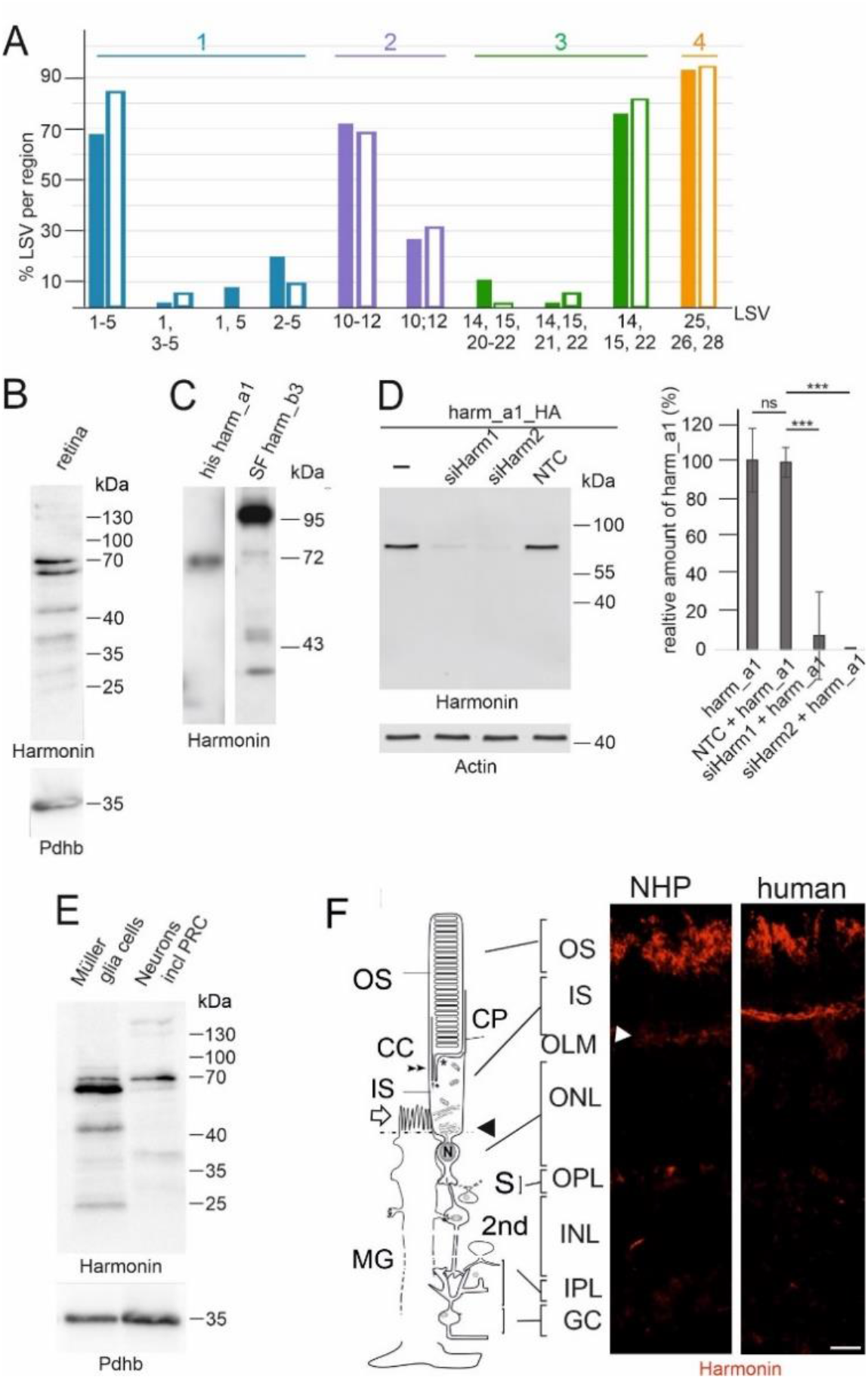
*USH1C*/harmonin expression in retinal cells. (**A**) Bulk RNA-seq of human retinal neurons (RNs) and Müller glial cells (MGCs). In RNs (filled bars) and MGCs (empty bars) *USH1C*/harmonin transcripts are detectable (**B**) Western blot analysis of harmonin protein expression in human retina with affinity-purified polyclonal antibodies against harmonin (H3). (**C**) His-tagged harmonin a1 and SF-tagged harmonin b3 were transiently expressed in HEK293T cells. Pan anti-harmonin H3 detected bands which co-migrate at the molecular weights of recombinant harmonin_a1 and harmonin_b3. Lower bands which may represent harmonin_c isoforms or degraded products of harmonin_a and/or b. (**D**) Western blot analysis to validate the specificity of the harmonin antibody. A strong harmonin band is detected in HEK293T cells transfected with HA-tagged harmonin (Harm_a1_HA) or co-transfected with Harm_a1_HA and control siRNA (NTC). Harmonin is not detected in cells co-transfected with HA-tagged harmonin_a1 (Harm_a1_HA) and siHarm, indicating the specificity of the harmonin antibody. P values: ***: < 0.001. (**E**) Western blot analysis of MGCs and RNs isolated from human retina. Both MGCs and RNs express various harmonin isoforms. (**F**) Localization of harmonin in retina sections. Indirect immunofluorescence labelling of harmonin in a longitudinal section through the retina, the outer segments (OS), the inner segments (IS) and the nuclei in the outer nuclear layer (ONL) and MGC of a non-human primate (NHP) and human retina, respectively. In addition to the prominent labelling of the outer limiting membrane (OLM, arrowhead), patchy harmonin staining was present in the layer of the photoreceptor OS. Faint staining was present in the IS, the outer plexiform layer (OPL), the inner plexiform layer (IPL) and ganglion cell layer (GCL). Scale bars: 10 µm.

In summary, our data demonstrate *USH1C* transcripts in both MGCs and RNs. Overall, the relative expression of the different LSVs between RNs and MGCs was quite similar and comparable to the quantification of the whole retinal samples.

### Harmonin protein expression and localization in human retinae

To assess harmonin protein expression in the human retina, we performed immunoblotting of protein lysates from human donor retinae using a harmonin antibody (H3), generated against a conserved *N-*terminal epitope (amino acids 1-89) of harmonin (21), suitable for detecting all five isoform groups of harmonin. Western blots demonstrated two major bands at a molecular mass close to 70 kDa (Figure 2B). The reference harmonin_a1 recombinantly expressed in HEK293T cells identified these bands as harmonin_a isoforms (Figure 2C), consistent with our RNA-seq data (Figure 1). In addition, we observed much less prominent higher molecular mass isoforms (∼130 kDa, ∼150 kDa) and two prominent lower molecular mass proteins of ∼45 and ∼38 kDa. These may represent harmonin homodimers (8) or either group c splice variants of harmonin (21, 36) or protein degradation products, respectively. The absence of any harmonin band in the immunoblots of the harmonin deficient tissue of a USH1C pig model (27) or of siRNA-mediated knockdowns of *USH1C*/harmonin in HEK293T cells further validated the specificity of our H3 antibody (Figure 2D). Immunoblots of fractions enriched for different cell types revealed harmonin protein variants in fractions of both MGCs and RNs (Figure 2E). Nevertheless, differences in the band pattern in the Western blots were striking: in the MGC, the slightly smaller band of harmonin_a (70 kDa) was much more pronounced than in the RN fraction and, in addition, further low molecular weight bands are found there that probably represent harmonin_c or protein degradation products.

To localize harmonin protein in the human and non-human primate (NHP) retina, we immunostained harmonin in retinal cryosections (Figure 2F). Fluorescence microscopy revealed intense harmonin immunofluorescence in the PRC outer segment (OS) layer and in the outer limiting membrane (OLM) of human and NHP retinae. In addition, we observed less intense immunostaining in the PRC inner segments, the outer and inner plexiform layers as well as the ganglion cell layer of human and NHP retinae.

### Localization of harmonin at the OLM junction complexes

OLM is characterized by heterotypic cell-cell adhesions between photoreceptor and MGC membranes and consists of proteins from both adherens and tight junctions (37). Immunofluorescence double staining of harmonin with β-catenin, a component of adherens junctions (38), with the tight junction molecule JAM-B (39), and with the actin-binding protein filamin A (40), respectively, showed partial co-localization with harmonin (Figure 3A). GST-pull downs of the GST-tagged β-catenin, JAM-B and filamin A from HEK293T cell lysates demonstrated that the interaction of the *C*-terminal tails of all three OLM junction proteins with His-tagged harmonin_a1 (Figure 3B, C, D). Reciprocal interaction pull-downs of GST-tagged PDZ domains revealed the binding of the *C*-terminal tail of β-catenin to the PDZ1 and PDZ3 domains of harmonin (Supplemental Figure S1A). Furthermore, yeast two-hybrid assays demonstrated the direct interactions of β-catenin, JAM-B, and filamin A with harmonin (data not shown). β-catenin presumably bind to the PDZ1 and PDZ3 domains of harmonin via a *C*-terminal type-I PDZ binding motif (DTDL) (41). In contrast, JAM-B contains a canonical type II PDZ targeting motif (FLV) at position 298-300 of its *C*-terminus, which is likely responsible for binding to harmonin (42).

**Figure 3.**
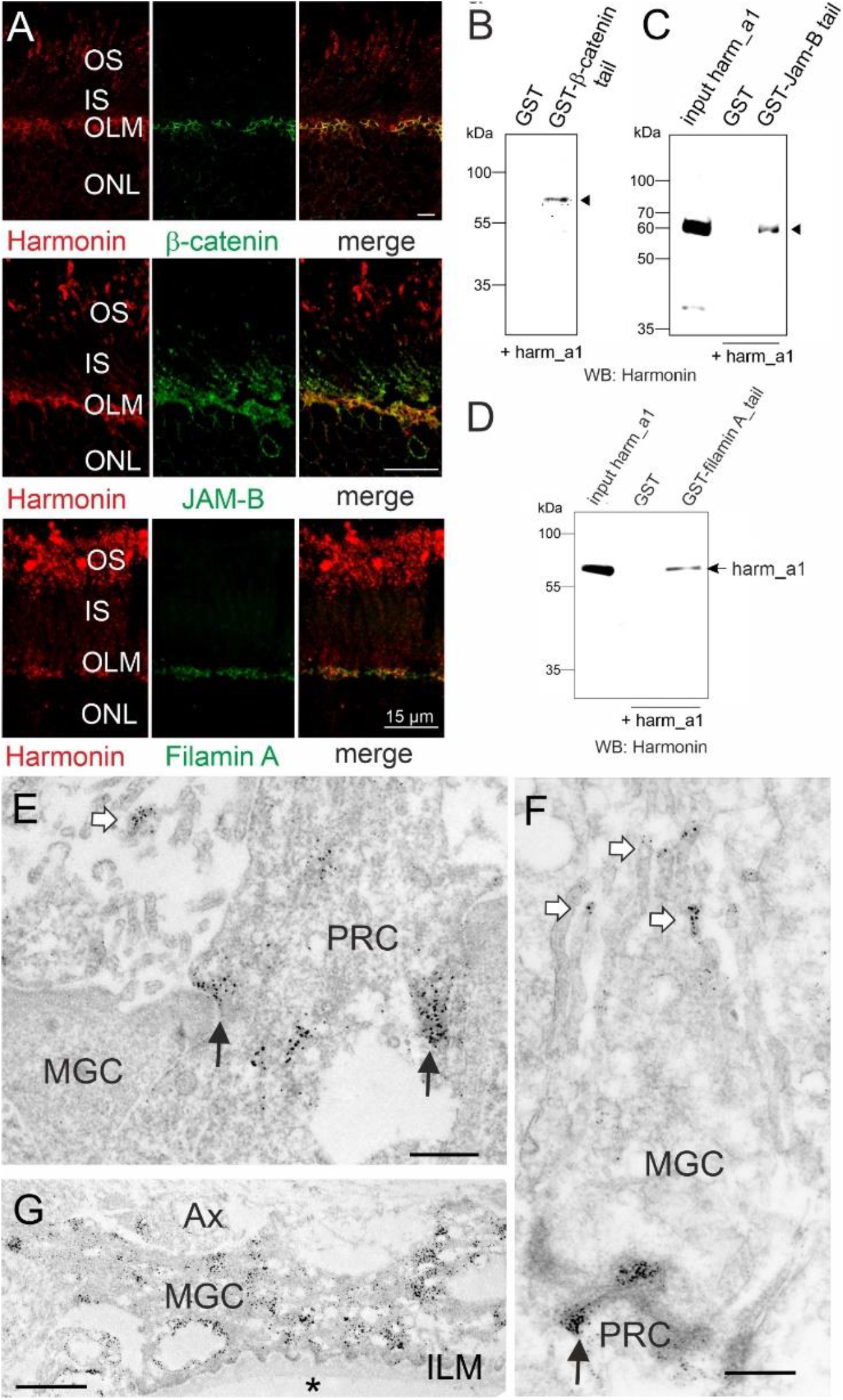
Subcellular harmonin localization at the outer limiting membrane of the human retina. (**A**) Indirect immunofluorescence double staining of harmonin (red) and the adhesion junction molecule β-catenin (green), the tight junction molecule JAM-B (green), or the actin-binding protein filamin A (green). Merged images demonstrate an overlap of harmonin staining with β-catenin, JAM-B and filamin A staining at the outer limiting membrane (OLM). (**B-D**) GST-pulldown demonstrates the interaction of harmonin_a1 (harm_a1) with the C-terminal tails of β-catenin, JAM-B, and filamin A, respectively. (**E-G**) Immunoelectron microscopy analysis of harmonin labelling in a longitudinal section through the OLM of a human retina. Harmonin labelling concentrated in the electron dense adhesion junctions between Müller glia cells (MGCs) and photoreceptor cells (PRC) in the OLM (black arrows), microvilli of MGCs (white arrows) and MGC endfeet at the inner limiting membrane (ILM) which contacts the vitreous (asterisk). OS, outer segment; IS, inner segment; ONL, outer limiting membrane; AX, axon; Scale bars: Harmonin/β-catenin: 5 µm; Harmonin/JAM-B, Harmonin/Filamin_A: 15 µm; D, E: 0.5 µm; F: 1 µm.

Immunoelectron microscopy of ultrathin sections through the OLM of the human retina localized harmonin to both sides of the OLM junctions in the photoreceptor cells and MGCs (Figure 3E, F). In addition, harmonin was labeled at the tips of microvilli (Figure 3E, F) and abundantly in the endfeet of the MGCs at the inner limiting membrane (Figure 3G).

### Differential expression of harmonin in rod and cone photoreceptor synapses

Next, we determined the localization of harmonin in photoreceptor synapses of the outer plexiform layer. We extended our analyses to NHP retinas because synaptic regions turned out not to be well preserved in human donor retinas, possibly due to postmortem changes. Immunofluorescence double staining revealed partial overlap of harmonin and RIBEYE, a marker of the synaptic ribbon (43) in photoreceptor synapses in both human and NHP sections (Figure 4A). Immunoelectron microscopy of NHP retinae allowed us to confirm the localization of harmonin at the photoreceptor synapses and uncovered dense harmonin labelling in pedicles of cones, whereas the labelling in rod spherules was much weaker (Figure 4B, C).

**Figure 4.**
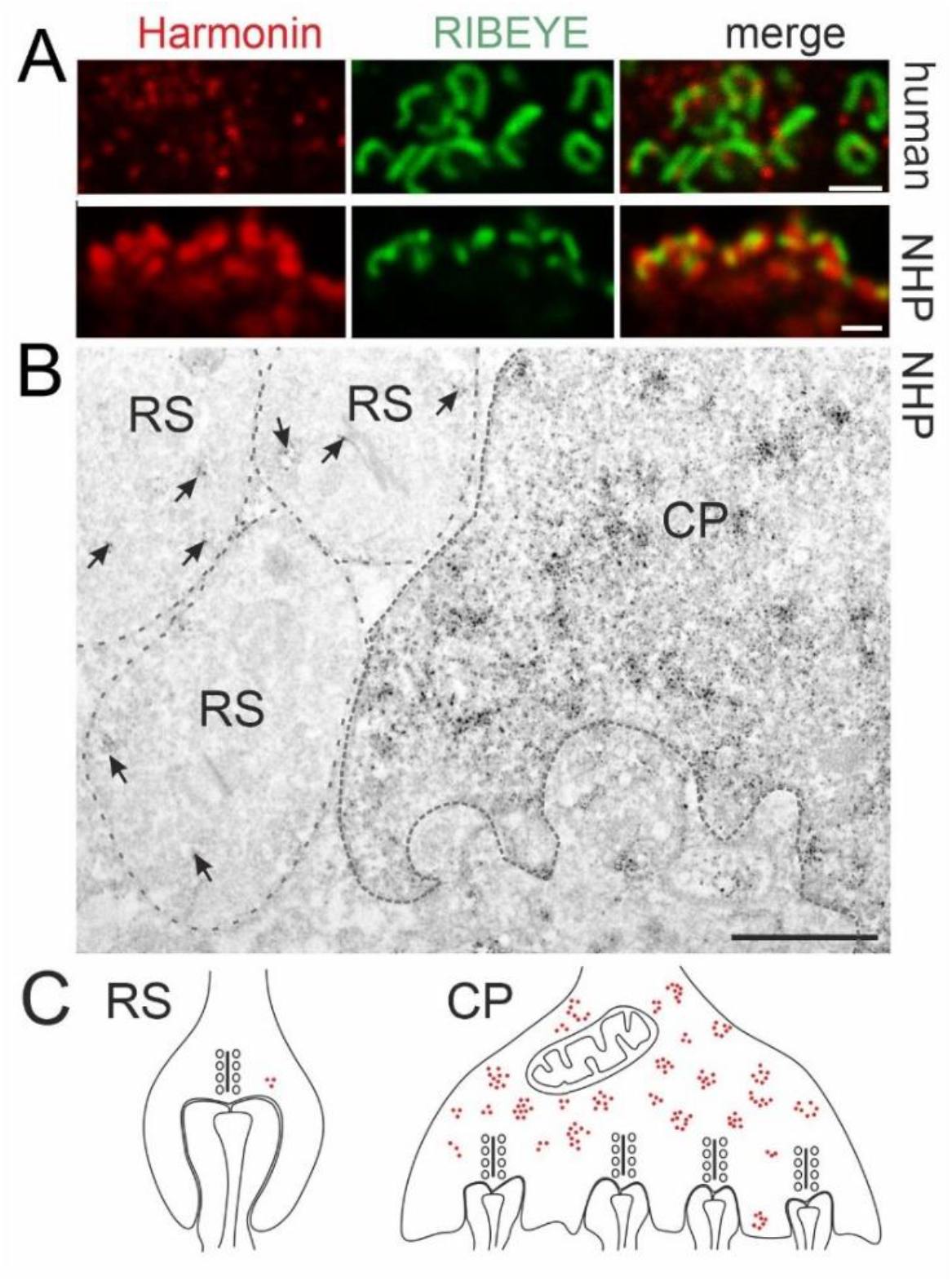
Harmonin localization at primate photoreceptor synapses. (**A**) Double immunofluorescence labelling of harmonin (red) and pre-synaptic protein RIBEYE (green) in human (upper panel) and non-human primate (NHP, lower panel) outer plexiform layer (OPL) synapses. Merged images revealed co-localization of harmonin and RIBEYE (yellow). (**B**) Immunoelectron microscopy analysis of harmonin in NHP photoreceptor synapses. Dense harmonin labelling was present in cone pedicles (CP), but only weak harmonin labelling (arrows) was observed in rod spherules (RS). (**C**) Schematic representation of presynaptic harmonin labelling in RS and CP. Scale bars: A, B: 1 µm; E: 12 µm

### Differential expression of harmonin in the outer segments of rod and cone PRCs

Immunofluorescence microscopy showed the presence of harmonin in the OS layer of the human retina (Figures. 2F, 5A). To discriminate between rod and cone OSs, we stained longitudinal cryosections through the human retina for harmonin and FITC-conjugated peanut agglutinin (PNA), a common marker for cones (44). Confocal analyses did not demonstrate a co-localization of harmonin and PNA (Figure 5B). This was confirmed by maximum projections of confocal planes of fluorescence images and z-sections of maximum projections. Furthermore, the overlay of fluorescent intensity plots of harmonin and FITC-PNA staining revealed apparent dissimilarities in their peaks (Figure 5C; low Pearson correlation coefficient values: Rr:0.18; and low Mander’s overlap coefficient values: M1:0.18; M2:0.19). In contrast, fluorescent intensity plots of harmonin and rod-specific arrestin-1 immunofluorescence staining showed apparent signal convergences (Figure 6C; high Pearson correlation coefficient values: Rr:0.59; and Mander’s overlap coefficient M1:0.57; M2:0.42) suggesting harmonin expression in rod, but not in cone OS (COS).

Immunoelectron microscopy of human PRCs confirmed our confocal data: harmonin labelling was detected in the OS of rods (ROS) but not of cones (Figure 5E-G). In the ROS, harmonin immunostaining was observed at the base where outer segment discs are formed *de novo* and throughout the outer segment along the cytoplasmic side of the disc membranes (Figure 5G). In addition, harmonin labelling was evident in the calyceal processes of PRCs, as previously described (23) (Figure 5G, asterisk). Western blots of purified OS fractions of porcine retinae confirmed the expression of harmonin in PRC OS (Figure 6A). Thus, our results show that harmonin is associated specifically with the disc membranes of ROS.

**Figure 5.**
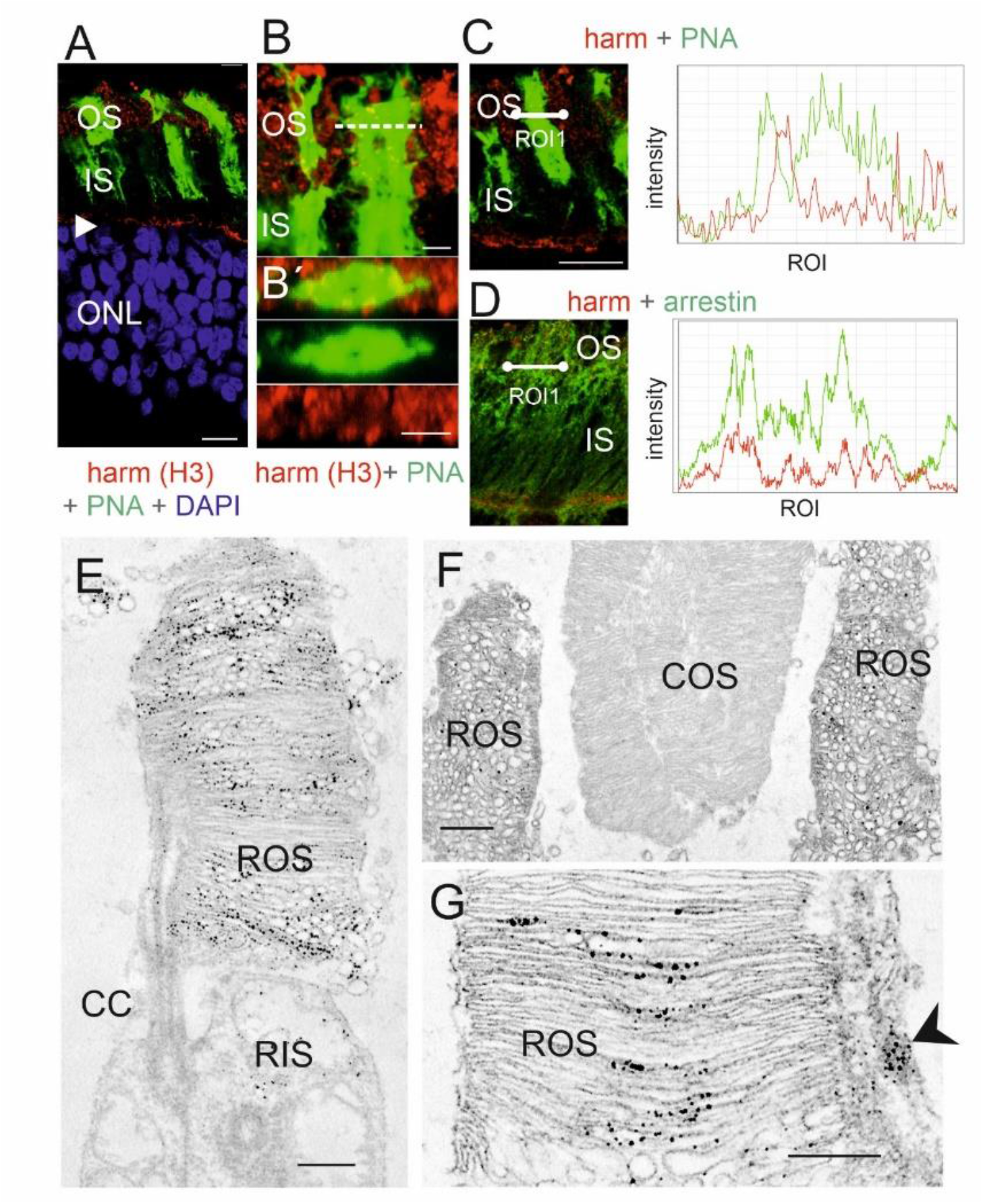
Harmonin in the outer segment of human photoreceptor cells. (**A**) Merged image of harmonin immunofluorescence (red) and fluorescent peanut agglutinin (PNA, green), a specific marker for the extracellular matrix sheath of cone photoreceptors in a longitudinal section through the photoreceptor layer, the outer segments (OS), the inner segments (IS) and the nuclei in the outer nuclear layer (ONL) of a human retina. In addition to the prominent labelling of the outer limiting membrane (arrowhead) patchy harmonin staining was present in the layer of the photoreceptor OS. (**B, B’**) Magnification of the OS region demonstrates no co-localization of harmonin and PNA (**B**’) Confocal x,y scan image of harmonin and PNA labelling in the photoreceptor layer of a human retina. y, z scan at the dotted line in B at higher magnification. (**C, D**) Double labelling of harmonin (red) with PNA (green) (**C**) and rod-specific arrestin (green) (**D**) in the photoreceptor layer of human retina. Corresponding fluorescence intensity profiles of regions of interest (ROI, white lines) at the right panel demonstrated no co-localization of harmonin and PNA, but co-localization of harmonin and arrestin indicating localization of harmonin in human rod outer segments but not cone outer segments. DAPI (blue): nuclear DNA. (**E-G**) Immunoelectron microscopy labelling of harmonin in a longitudinal section through human retinae. (**E**) In human rod PRCs, harmonin was labelled in rod outer segments (ROS) and was barely detected in rod inner segments (RIS). (**F**) Harmonin was labelled in ROS, but not in outer segments of cones (COS). (**G**) Harmonin is detected in ROS and calyceal processes (arrowhead). CC: connecting cilium. Scale bars: A: 10 µm, B: 2.5 µm, B’C, D: 5 µm, E, G: 0.5 µm, F: 1 µm

**Figure 6.**
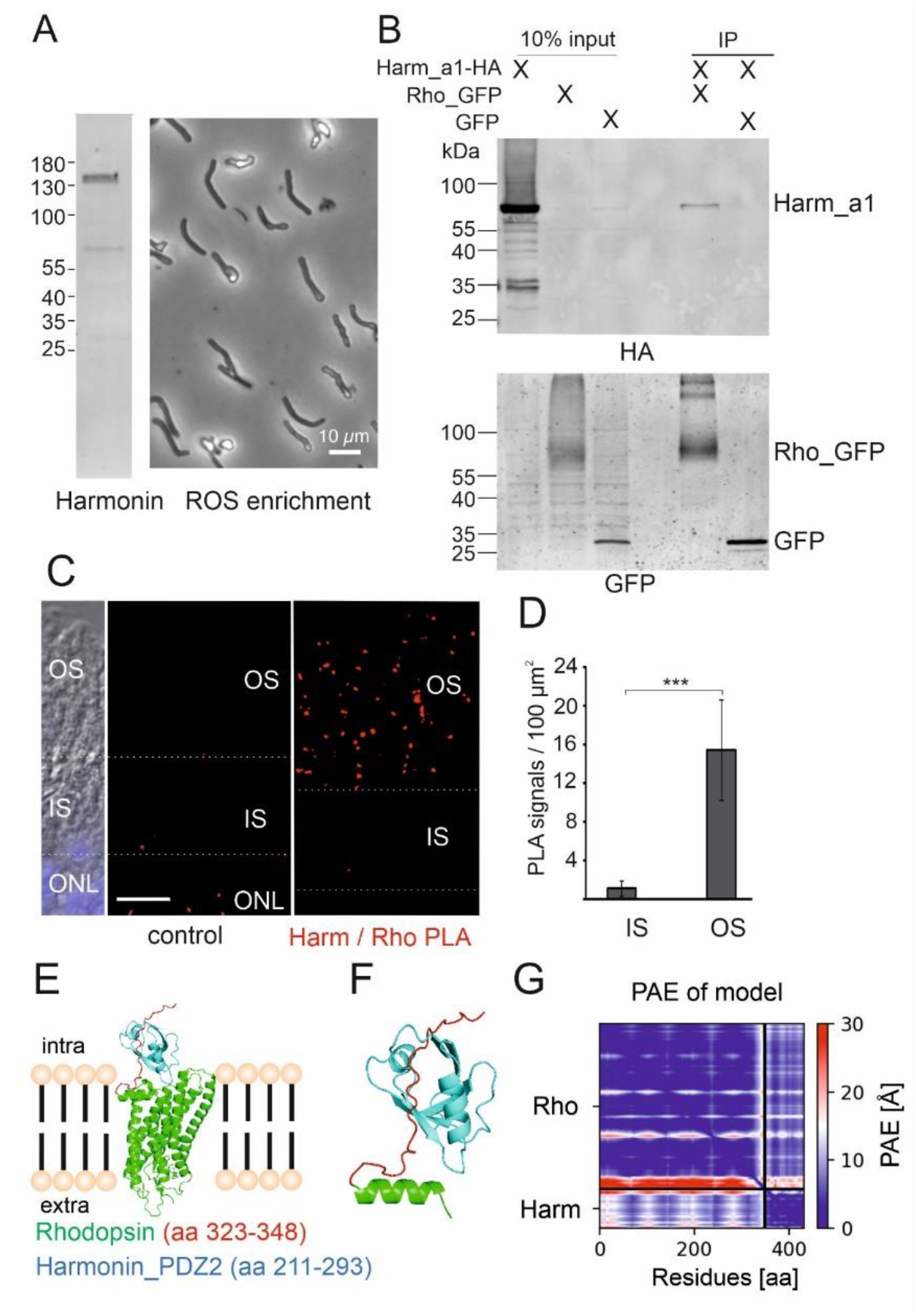
Harmonin is expressed in rod outer segments and interacts with rhodopsin. (**A**) Harmonin is expressed in porcine photoreceptor outer segments. Left panel: Western blot analysis of the isolated porcine ROS enriched fraction from density gradients revealed harmonin expression. Right panel: phase contrast picture of material used for the Western blot. (**B**) GFP-Trap® demonstrates interaction between harmonin a1 (Harm_a1) and opsin-GFP (Rho_GFP) in transfected HEK293T cells. Harmonin_a1 was precipitated by immobilized opsin-GFP, but not by GFP alone (replicates, n=3) (**C**) Proximity ligation assay (PLA) of harmonin (Harm) and rhodopsin in longitudinal cryosections through a human retina. PLA signals (red dots) indicate the interaction of harmonin (Harm) and rhodopsin in OS of human PRCs *in situ*. For quantitative analysis the inner segment/outer segment (IS/OS) borders were defined based on DIC images. Nuclear DAPI staining (blue) was used to define the outer nuclear layer (ONL). ImageJ was adopted to define the different retina layers: OS and IS of PRCs, and ONL (white dashed lines). As control, PLA was performed anti-opsin antibodies and oligonucleotide-labelled antibodies, where almost no PLA signals were found. (**D**) Quantification of PLA signals in the OS and IS of PRCs. PLA signals were counted automatically in the different compartments and signals in controls were subtracted in three different samples. The number of PLA signals in OS were also significantly higher when compared to signals in IS. Scale bar: 10 µm. (**E-G**) Structure of the human rhodopsin-harmonin_a1_PDZ2 protein complex predicted by AlphaFold2. (**E**) Structure of full-length rhodopsin (Rho) (green) and harmonin (Harm)_PDZ2 (aa 211-293, blue). AlphaFold2 predicts binding of Harm_PDZ2 to intracellular *C-*terminus of Rho (aa323-348), enlarged in (**F**). (**G**) Predicted Alignment Error (PAE) for the modelled structure is very low for the predicted complex Harm_PDZ2-Rho (lower left and upper right rectangle) indicating in high confidence of the protein complex. Heat map illustrates amino acid distances in Å (further explanation, see Supplemental Figure S2).

### Harmonin interacts with rhodopsin

In the microvilli of the rhabdomes of *Drosophila* photoreceptors, functionally homologous to the outer segment discs of vertebrate PRCs, rhodopsin binds to the PDZ domain-containing scaffold protein INAD (45). To test whether the PDZ-protein harmonin also interacts with human rhodopsin, we performed *in vitro* co-immunoprecipitations assays with protein lysates from HEK293T cells expressing GFP-rhodopsin and HA-tagged harmonin or RFP-harmonin, respectively. Western blots of the recovered proteins revealed that harmonin_a1 co-immunoprecipitated with GFP-rhodopsin, but not in controls with GFP alone (Figure 6B; Supplemental Figure S1B).

To examine a putative interaction of harmonin and rhodopsin in the human retina, we performed *in situ* proximity ligation assays (PLA) applying anti-harmonin (H3) and monoclonal antibodies to rhodopsin in retinal cryosections (Figure 6C). We observed positive PLA signals in the photoreceptor layer, predominately in the OS layer and a significant smaller numbers of signals in the inner segment (Figure 6C,D). In contrast, almost no positive PLA signals were found in the negative controls (Figure 6C).

*In silico* modelling of molecular interaction of harmonin and rhodopsin applying AlphaFold2/Multimer (46, 47) predicted a complex of harmonin_PDZ2 and cytoplasmic *C-* terminus of rhodopsin with high confidence of the predicted alignment error (PAE) (Fig. 6E). In contrast other domains of harmonin were not predicted to bind to rhodopsin (Supplemental Material Figure S2B-G). Moreover, modelling of the harmonin_PDZ2 of with the adrenergic receptor α-A1 did also not predict confident complex formation (Supplemental Figure S2H) further supporting the specificity of the harmonin_PDZ2-rhodopsin binding.

In summary, harmonin localization in ROS shown by complementary methods of light and electron microscopy, paired with the interaction of harmonin and rhodopsin *in vitro*, the close proximity of both proteins in the OS layer demonstrated by *in situ* PLAs as well as AlphaFold2 *in silico* predictions provide first evidence for the interaction of rhodopsin and harmonin in ROS.

### Retinal phenotype of USH1C patients

Two male siblings with confirmed mutations in *USH1C* (c.91C>T;p.(R31*), c.238dupC;p.(Arg80Profs*69)) were clinically examined at the age of 35 and 47 years, respectively (Figure 7A-E; Supplemental Figure S3). Optical coherence tomography (OCT) analysis of the retina of 35-year-old male patient revealed atrophy and thinning of the photoreceptor outer/inner segments and of the outer nuclear layer up to the fovea. Outer retinal atrophy progressed to constriction of the visual field at around 5 degrees and a hyperfluorescent perifoveal ring on the fundus autofluorescence imaging (Figure 7B). Additionally, epiretinal gliosis observed in OCT is a common complication of PRC degeneration which also was a hallmark in the USH1C pig model (27). The posterior eye pole showed typical bone spicules in the mid periphery and periphery (Fig. 7A). Still, despite progressive peripheral retinal degeneration at 35 years, the central vision of the patient was relatively well preserved with visual acuity of 20/25 in both eyes and normal colour perception (Figure 7E).

**Figure 7.**
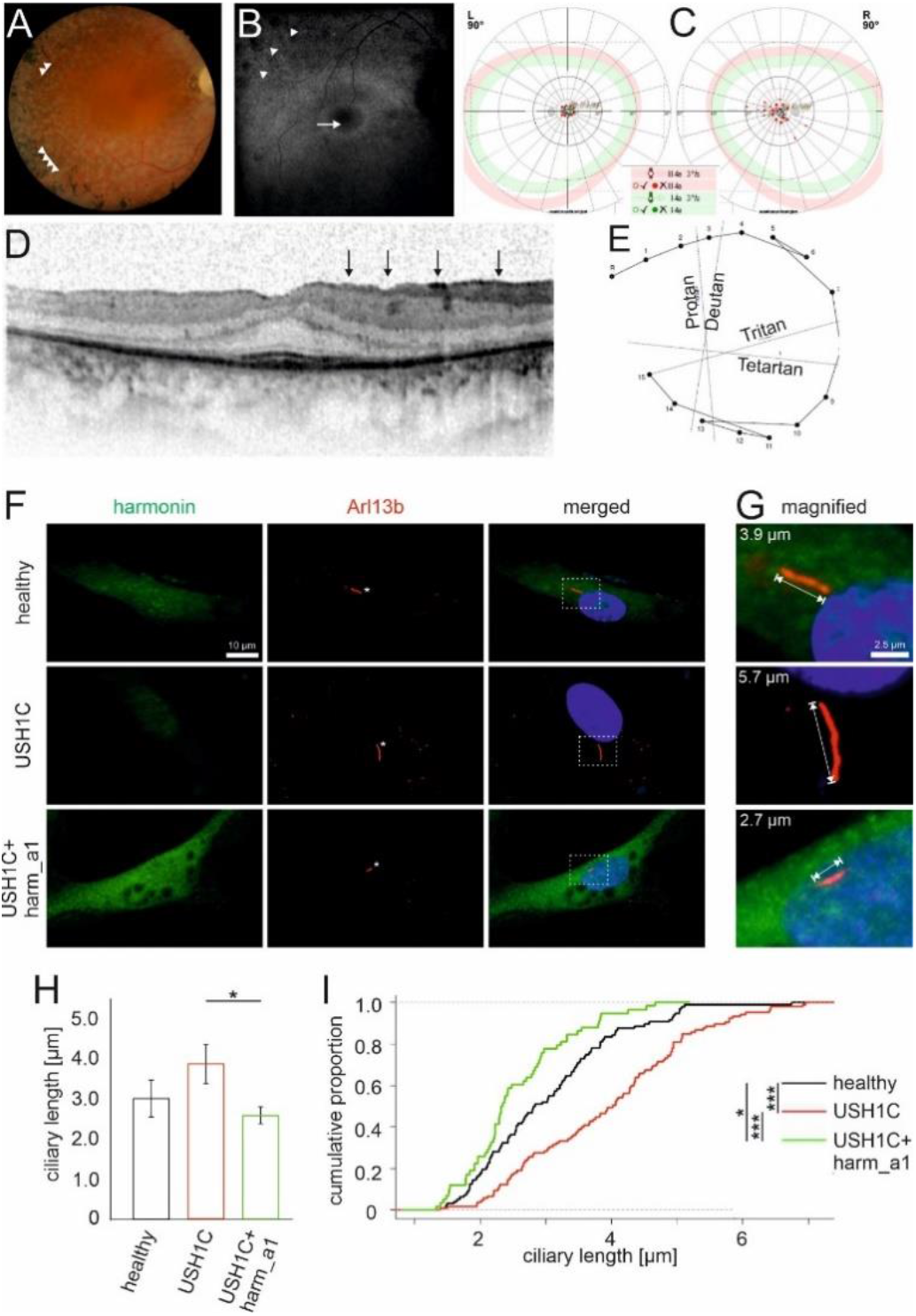
Retinal phenotype of an USH1C patient and ciliary phenotype of USH1C patient-derived cells. **(A-E)** Clinical findings of the retinal phenotype of a 35-year old male with confirmed mutations in USH1C (c.91C>T;p.(R31*), c.238dupC;p.(Arg80Profs*69)). (**A**) Fundus photography showed bone spiculas in the mid peripheral and peripheral retina (arrowheads), attenuated retinal vessels, waxy pallor optic disc, and white spots of retinal pigment epithelium atrophy. (**B**) Fundus autofluorescence imaging displayed a hyperfluorescence ring around the fovea (arrow) and a disrupted hypofluorescence in the mid- and far periphery of the retina (arrowheads) corresponding to the outer retinal atrophy. (**C**) Kinetic visual field (90°): Concentric constriction for III4e and I4e markers with central preserved area. (**D**) Optical coherence tomography showed epiretinal gliosis (marked as black vertical arrows), as well as gradual IS/OS loss up to the fovea. Only the foveal region displays a normal retinal structure with preserved photoreceptors and a central macula thickness 269 µm. (**E**) Lanthony color test showed normal color perception. **(F-I)** Primary ciliary phenotype of USH1C patient-derived cells and rescue by harmonin_a1. (**F, G**) Immunofluorescence of harmonin (green) and ciliary marker Arl13b (red) in fibroblasts from a healthy donor (healthy), the clinically examined USH1C^R80Pfs*69/R31*^-patient (USH1C) and USH1C-fibroblasts transfected with harmonin_a1 (USH1C+harm_a1). (**F**) In healthy donor cells and harmonin_a1 transfected USH1C^R80Pfs*69/R31*^ cells, harmonin is detectable. In untransfected USH1C^R80Pfs*69/R31*^ cells, harmonin staining is barely visible. (**G-I**) Ciliary length measurements revealed longer cilia in USH1C patient-derived cells compared to control cells (healthy) and harmonin_a1 transfected USH1C cells. (**H**) Quantitative analysis of primary ciliary length reveals significant decrease of ciliary length in USH1C+harm_a1 fibroblasts towards the ciliary length of healthy donors. (**I**) Cumulative analysis of ciliary length and number of ciliated cells in fibroblasts of healthy donors (healthy), USH1C^R80Pfs*69/R31*^ (USH1C), and USH1C^R80Pfs*69/R31*^ fibroblasts transfected with harmonin a1 (USH1C+harm_a1) using R. Two-tailed Student’s t test, *p≤0.05, **p≤0.01. Cells analysed: healthy: n=96; USH1C: n=105; USH1C+harm_a1: 58, in three independent experiments. Scale bar: F: 10 µm, G: 2.5 µm.

### Rescue of the ciliary phenotype in patient-derived fibroblasts

The photoreceptor outer segment resembles a highly modified primary sensory cilium, which has been shown to be impaired in retinal ciliopathies such as USH (4, 5, 48). We analysed primary cilia of dermal fibroblasts from one of the clinically characterized USH1C^R80Pfs*69/R31*^ patient. After induction of ciliogenesis (3) primary cilia of USH1C patient fibroblasts were significantly longer compared to the primary cilia of healthy donor fibroblasts (Figure 7F-I). Strikingly, this ciliary phenotype of USH1C^R80Pfs*69/R31*^ patient fibroblasts was rescued by re-expression of the harmonin_a1 (Figure 7F,I). These data demonstrated that USH1C patient cells can serve potential cellular models for evaluating therapies for USH1C and that the harmonin_a1 isoform is a therapeutic active isoform, able to revert the USH1C phenotype to normal.

## Discussion

Alternative splicing regulates and increases the diversity of the transcriptome and proteome. Tissue- and cell specific alternative splicing frequently occurs, previously also predicted for *USH1C* (36, 49). In comparison to other tissues, the neuronal retina displays the highest level of alternative splicing events (50, 51). Splicing programs are essential for retinal function and maintenance, and are regulated in an extremely complex way by retina-specific splicing factors (52-54). Here, we demonstrate that *USH1C* is extensively alternatively spliced in the human retina. We do not expect all variants observed to be translated into protein due to pre-translational processing of mRNAs (55). Nevertheless, we assume that several *USH1C*/harmonin isoforms are expressed, potentially with distinct patterns in the various human retinal cell types providing cell-specific functions as indicated in the cochlear and vestibular hair cells of the inner ear (49).

Harmonin is a scaffold protein that is modularly composed of well-defined domains and motifs, such as the *N*-terminal HHD, PDZs, CCs and a PBM (Fig. 1A,G), which all participate in protein-protein interactions (8, 16). Identification of LSVs that include specific domains suggests that alternative splicing may affect binding of harmonin to target molecules in human retinal cells. For example, alternative splicing of exon 2 in region 1 is predicted to alter the globular *N*-domain, termed HHD, and influence the binding affinity of harmonin PDZ1 to target proteins, such as VLGR1/ADGRV1 (USH2C), USH2A and SANS (USH1G) (14, 15). This hypothesis is supported by our recent results indicating reduced binding affinity of target proteins to harmonin lacking parts of the *N*-terminal HHD (Sturm, 2020). Furthermore, splicing out of exons 16-21 in region 3 would lead to the loss of CC2 and PST domains, which should impact oligomerization of harmonin molecules and loss of actin filament bundling properties, respectively (17).

Differential binding of proteins to the numerous harmonin splice variants may imply differences in harmonin function in various retinal cell types. Accordingly, the control and regulation of alternative splicing of *USH1C*/harmonin in the retina is of importance. Previous studies have identified Musashi proteins (MSI1 and MSI2) as important regulators of alternative splicing in the retina (53, 54, 56). However, *USH1C*/harmonin has not been identified as a target for the splicing control machinery of Musashi proteins. Recently, we were able to show that the USHG1 protein SANS interacts with core components of the spliceosome and regulates splicing including constitutive and alternative splicing of *USH1C*/harmonin (57). In particular, the latter study shows that SANS causes the retention of exon 11, thereby leading to alternative expression of the human harmonin_a1 and _a4 transcripts. Further studies are needed to understand the interplay and interference of the two USH1 proteins, SANS and harmonin, in the retina at two different levels: in protein complexes (SANS-SAM/PBM binding to the *N*-terminal HHD/PDZ1 and PDZ3 of harmonin), possibly related to transport processes (14, 16, 58), and during splicing where SANS presumably modulate *USH1C*/harmonin pre-mRNA (57).

The assignment of LSVs to *USH1C*/harmonin domains allowed us to quantify the expression of the harmonin classes in the human retina (Figure 1G). This analysis demonstrates that harmonin_a is by far the most abundant class expressed in the human retina in concordance with our results by semi- and quantitative RT-PCR (Figure 1B-C) and immunoblotting analyses (Figures 2B-E) of donor retinae. Our data also indicate that the *USH1C*/harmonin_a1 transcript is more frequently expressed than *USH1C*/harmonin_a4 in the retina. Based on these findings, we concluded that harmonin_a1 is the most promising variant for gene replacement therapies. The rescue of the pathogenic phenotype in primary cilia of fibroblasts of USH1C patient after addition of harmonin_a1 further supports this conclusion (Figure 7).

In contrast to the prevailing hypothesis of *USH1C*/harmonin expression in PRCs (21, 23), our results reveal *USH1C*/harmonin expression in RNs (mainly PRCs) and MGCs of the human retina. Almost equal levels of *USH1C* expression were detected in RNs and MGCs (Figure 2A) by bulk RNA-seq of donor retinae. Of note, we did not observe substantial differences in LSV expression levels in RNs and MGCs. Our findings don’t support a recently published scRNA-seq study indicating *USH1C* expression almost exclusively restricted to MGCs in the human retina (29). The limited sequencing depth in scRNA-seq likely accounts for this discrepancy. In addition, scRNA-seq requires dissociation of retinal cells prior to RNA-seq library preparation, which can have a huge impact on the gene expression profile (59). Our results from expression profiling are confirmed by immunoblotting and immunocytochemical studies by light and electron microscopy that demonstrate expression and subcellular localization of *USH1C*/harmonin in MGCs as well as PRCs.

The localization of harmonin in distinct subcellular compartments of human MGCs may provide insights into harmonin function. Immunoelectron microscopy revealed the localization of harmonin to the tips of the microvillar processes formed by their apical membrane of MGCs which project into the subretinal space surrounding the PRC inner segments (Figure 3E, F), consistent with localization of harmonin in both the tip-link complex of stereocilia (which resemble modified microvilli) of auditory hair cells (60, 61) and the very similar tip-link complex of the brush border microvilli of intestinal enterocytes (62). In contrast to tip-link complex of stereocilia, in which harmonin interacts with other USH proteins, in the brush border microvilli tip-link, harmonin forms a stable ternary complex with ANKS4B and MYO7B for anchoring specific cadherins (62, 63). A morphologic similarity between the microvilli of MGCs and the brush border microvilli is evident. Given that the absence of tip-link molecules in brush border microvilli leads to structural perturbations linked to ineffective epithelial transport (64), we suggest that harmonin may contribute to the structural arrangement of the microvilli populating the apical MGC membrane, being essential for their physiological function (31). Here we provide first evidence for a tip-link complex in the microvilli of MGCs with a possible scaffolding function of harmonin. Additionally, we identified abundant harmonin expression in conical endfeet at the other pole of the MGCs adjacent to the inner limiting membrane and vitreous humor (Figure 3G). Harmonin may be integrated into protein networks that mechanically stabilize the endfeet at the MGC base and may support anchoring of MGCs at the inner limiting membrane.

Furthermore, we show a prominent submembranous localization of harmonin in MGCs and PRCs associated with the specialized heterotypic adhesion junctions of the OLM (Figure 3). Partial co-localization of harmonin and β-catenin, a linker of cadherins to actin filaments, and the transmembrane protein JAM-B, together with the binding of the cytoplasmic tails of both proteins to harmonin strongly suggest that harmonin provides the molecular scaffold to anchor both proteins in the submembranous cytoplasm at the OLM junctions. Harmonin perhaps contributes to the linkage of the OLM junction complexes to the actin cytoskeleton. This is supported by the observed co-localization and binding of harmonin to the actin-binding protein filamin A, which has been described as an important component of the actin cytoskeleton in barrier-forming cell-cell adhesion complexes (87). Since harmonin is present in both adhesion and tight junction complexes, its deficiency may disrupt the structural integrity of the retina (adhesion component) and/or increase the permeability (tight junctional component) of the retinal barrier. In pathological conditions, the disruption of the retinal barrier function of the OLM would contribute to fluid accumulation in the macula, clinically manifesting as macular edema, which can be detected in many hereditary retinal degenerative diseases, including USH (65). In concordance, OCT of the two USH1C patients, reported here, not only show thinning of the PRC layer but also reveal macular alterations (Figure 7A-E, Supplementary Figure S2). Further studies are required to elucidate whether macular lesions are caused by impaired leaky OLM adhesion junctions or by other processes, *e*.*g*., MGC gliosis or inflammation.

As in MGCs, we also observed the localization of harmonin in distinct subcellular compartments of cone and rod PRCs, namely in submembranous cytoplasm of the basal inner segment at the heterotypic OLM junctions (see above), outer segments, calyceal processes, and ribbon synapses. Previous studies indicate a multifaceted presynaptic role of harmonin in ribbon synapses, regulating L-type Ca^2+^ channels and neurotransmitter exocytosis in inner ear hair cells (18, 19, 66). Though harmonin is detected in the ribbon synapses of both PRC types, our immunoelectron microscopy data show that harmonin is more abundant in the synaptic pedicles of cones than in rod presynaptic terminals. This is consistent with the previously reported role of harmonin in synaptic maturation in cone pedicles of zebrafish (26) and with recent findings in our USH1C knock-in pig model indicating that the harmonin deficiency alters the width of cone synaptic pedicles (27). Notably, earlier studies have suggested selective modulation of pre-synaptic L-type calcium currents in cones to broaden the dynamic range of synaptic transfer by controlling transmitter release (67). Our data therefore support a putative role of harmonin contributing to this process.

The localization of harmonin in the calyceal processes of PRCs is consistent with previous reports (23). There is evidence that cadherin 23 (USH1D) and protocadherin 15 (USH1F) link the membrane of the calyceal processes to the plasma membrane of the outer segment via their long extracellular domains (28, 68, 69). In this scenario, harmonin is thought to anchor both cadherins in the cytoplasm of the calyceal processes mediated by binding of their *C*-terminal PBMs to harmonin’s PDZ2. More strikingly, we consistently found abundant harmonin expression in the outer segment of rod PRCs, whereas no harmonin is present in cone outer segment. There are obvious differences in disc morphogenesis and disc stacking between rod and cone PRCs. In contrast to cones, rod coin-roll like stacked disc membranes become rimmed by the plasma membrane during disc neogenesis at the base of the outer segment (70, 71). The localization of harmonin at the base of the outer segment and its ability to bind actin are consistent with its involvement in disc biogenesis, with the actin cytoskeleton playing a key role (72, 73).

Harmonin lines up along the disc membranes of the entire rod outer segment indicating a role in disc stacking. There is evidence for a network of cytoplasmic membrane-membrane tethers and/or spacers localized between the outer segment discs, but their molecular identity remains to be elusive (71). Interestingly, harmonin fulfills the criteria of such spacers: as a scaffold protein, harmonin facilitates protein networks, forms homomers, binds to membranes, directly to phospholipids, and interacts with transmembrane proteins (8, 74) such as rhodopsin, as shown herein. Our data on the interaction between harmonin and rhodopsin also suggest a role for harmonin in anchoring rhodopsin in disks by binding to its cytoplasmic *C*-terminus. Such roles of harmonin in disc morphogenesis and stacking are in line with our recent findings on the altered disc architecture of ROS in the USH1C pig model, revealing vertically oriented membrane discs, disk stacks interrupted by interstitial gaps and vesicle-like structures present at the outer segment base of rods in the absence of harmonin (27). Nevertheless, further studies are necessary to test and validate harmonin functions in the rod outer segment.

In conclusion, we show that the scaffold protein harmonin is expressed in the human retina in form of numerous splice variants, which localize in different subcellular compartments of PRCs and in MGCs. Defects in *USH1C*/harmonin would likely impair its various functions in distinct subcellular compartments of PRCs and MGCs. We hypothesize that these cellular dysfunctions have a cumulative effect, leading to progressive though slow retinal degeneration and vision impairment characteristic of USH1C patients. Understanding how these cellular defects interfere, potentially amplify, and accumulate will provide new clues to cellular pathophysiology, elucidating potential targets for treatment and cure of USH1C patients. Although we cannot provide a fully elaborated therapeutic concept, we provide evidence that harmonin_a1 is the most promising splice variant for gene supplementation therapy and also identify both PRCs and MGCs as targets to treat vision loss in USH1C-patients.

## Material & Methods

### Human Subjects

Procedures adhered to the Declaration of Helsinki and were approved by the institutional review boards. Written informed consent was obtained from patients and probands. All analyses of human retinae were performed using *post-mortem* anonymized samples (Supplemental Table S1). Donors had no known history of retinal disease and were either obtained from the Department of Ophthalmology, University Medical Center Mainz, Germany, the University of Utah, USA or from the National Disease Research Interchange (NDRI, Philadelphia, USA) and conformed to the tenets of the Declaration of Helsinki.

### Clinical examination

Two sibling patients genetically diagnosed for USH1 with biallelic *USH1C* mutations (c.91C>T;p.(R31*), c.238dupC;p.(Arg80Profs*69) were examined clinically at the Center for Ophthalmology, University of Tübingen, including high-resolution retinal imaging and multimodal functional diagnostics. The clinical examination included a detailed medical history, best corrected visual acuity (BCVA) testing, color perception evaluation using the Lanthony test, kinetic perimetry (Octopus 900; Haag-Streit International, Germany), slit-lamp examination, fundus examination in mydriasis with color fundus photography and 30 degrees fundus autofluorescence, as well as spectral domain optical coherence tomography (OCT) (Heidelberg Engineering GmbH, Germany).

### Non-human primates (NHP)

Eyes from three adult unaffected macaques (*Macaque mulatta*) were obtained from the German Primate Center (DPZ) where they were sacrificed as controls in other unrelated experiments.

### Transcriptome analysis

RNA isolation from donor retinae was performed with “RNeasy Mini Kit” (Qiagen, Hilden, Germany), according to company’s instruction. To increase the RNA yield, elution was performed twice. Libraries were generated with Illumina’s TruSeq Stranded mRNA Library Prep Kit and sequenced using Illumina HiSeq2500 sequencing platform as stranded, paired-end with a length of 151-nucleotides (mRNA datasets) (*IMSB*, Mainz, Germany). Using the “best practice” pipeline from NEI commons using Trimmomatic (75), STAR (76), and Kallisto (77) (https://neicommons.nei.nih.gov/#/howDataAnalyzed), we aligned our sample to the Ensembl transcriptome version 89. 81.4% of the 89.2 million reads derived from Donor 269-09 (Supplemental Table S1) were aligned to the respective genome reference, out of which ∼96% aligned uniquely. The mapped paired-end read length was 289.4 bp (of 302 bp) with a mismatch rate of 0.2%. Previously published RNA-seq samples (35) were downloaded as BAM-files from EBI’s ArrayExpress (E-MTAB-4377), reverted to FASTQ-files using SAMTOOLS (78) and processed the same way as mentioned above. For visualization we used the Integrative Genomic Viewer (IGV, https://www.broadinstitute.org/igv/) (79, 80).

### Reverse transcription, polymerase chain reactions, subcloning, and identification of LSVs

The RNA isolation was performed using “TRIzol” (Invitrogen, USA) according to company instructions. The reverse transcription (RT-PCR) was performed with “Superscript II Reverse Transcriptase” (Invitrogen), according to company instructions. The polymerase chain reaction (PCR) was performed with “Taq DNA Polymerase with Standard Taq Buffer” (NEB) according to company’s instructions. qPCR was performed on CFX96 real-time system (Bio-Rad) using the SYBRGreen iTAQ according to manufacturer’s instructions. Primers based on human *USH1C* sequences (http://www.ncbi.nlm.nih.gov/gene/10083) are listed in Supplemental Table S3.

### MACS-sorting of human MGCs and RNs

Retinal cell types were enriched using magnetic-activated cell sorting (MACS) as described previously (81).

### Bulk RNA-sequencing on purified retinal cell populations

Total RNA was isolated from cell pellets after immunoseparation using the PureLink^®^ RNA Micro Scale Kit (Thermo Fisher Scientific, Schwerte, Germany). RNA integrity validation and quantification was performed using the Agilent RNA 6000 Pico chip analysis according to the manufacturer’s instructions (Agilent Technologies). Enrichment of mRNA and library preparation (Nextera XT, Clontech), library quantification (KAPA Library Quantification Kit Illumina, Kapa Biosystems, Inc., USA) as well as sequencing on an Illumina platform (NextSeq 500 High Output Kit v2; 150 cycles) were performed at the service facility of the KFB Center of Excellence for Fluorescent Bioanalytics (Regensburg, Germany; www.kfb-regensburg.de). After de-multiplexing, at least 20 million reads per sample were detected. Quality control (QC) of the reads and quantification of transcript abundance was performed applying STAR (76), *cutadapt* was used to remove adapter sequences and several QC measures were queried with *fastqc*. Trimmed reads were aligned to the reference genome/transcriptome (*mm10*) applying STAR and transcript abundance was estimated with RSEM (82) expressed as **T**ranscripts **P**er kilobase **M**illion (TPM).

### Western blot analyses

The immunoblots were performed as previously described (83). Briefly, one-fourth of a human retinae was placed in 125 µl HGNT-buffer, sonicated 3 times for 5 s and mixed with SDS-PAGE sample buffer (62.5 mM Tris-HCL, pH 6.8; 10% glycerol, 2% SDS, 5% mercaptoethanol, 1 mM EDTA and 0.025% bromphenol blue). As a molecular marker PageRuler™ Prestained Protein Ladder ranging from 11 to 170 kDa was used (Fermentas). 30 µg retina protein extracts were separated on 8% polyacrylamide gels and blotted onto PVDF transfer membranes (Millipore) followed by blocking with non-fat dried milk (Applichem). Cell pellets of enriched cell populations from retinal punches (6 mm) were dissolved in reducing Laemmli sample buffer, denatured and sonicated. Samples were separated using a 12% SDS-PAGE. Detection was performed either by a chemiluminescence detection system (WesternSure PREMIUM Chemiluminescent Substrate, LI-COR) or using the Odyssey infra-red imaging system (LI-COR Biosciences).

### Antibodies and fluorescent dyes

Primary antibodies used: Anti-Arl13b (ab136648, Abcam), anti-JAM-B (SAB2501282, Sigma-Aldrich), anti-β-catenin (cs-7963, Santa Cruz), anti-filamin A (67133-1-ig, Proteintech) and anti-arrestin (SCT128, Santa Cruz Biotechnology), anti-his (27-4710-01, Amersham™), anti-actin (MA5-11869, Thermo Fisher Scientific), anti-GFP (gift from Clay Smith), anti-PDHB (ab155996, Abcam) and anti-RFP (6G6, Chromotek). The antibody against harmonin (H3) was previously described (21). Monoclonal antibodies (mAbs) against bovine rod opsin B6-30a1, K16-155, and R2-15 were applied as previously described (84, 85). Subcellular markers were RIBEYE (612044, BD Bioscience), fluorescein-labelled lectin peanut agglutinin (FITC-PNA) and 4’,6-diamidino-2-phenylindole (DAPI) (Sigma-Aldrich). Secondary antibodies for immunofluorescence and Western blot analysis were conjugated to Alexa 568 and Alexa 488 (Molecular Probes) or coupled to horseradish peroxidase (ECL Plus Western Blotting Detection System, GE Healthcare), respectively.

### Immunofluorescence microscopy

After dissection human retinae and NHP retinae were cryofixed 11.5 to 31 hours *postmortem* or directly, respectively, cryofixed in melting isopentane, cryosectioned and stained immunofluorescence as previously described (86, 87). Immunofluorescence microscopy was performed with a Leica SP5 confocal laser scanning microscope (Leica microsystems) or a Leica DM6000B deconvolution microscope (Leica). Images were processed with Adobe Photoshop CS (Adobe Systems). Colocalization analysis were performed with the ImageJ (http://rsbweb.nih.gov/ij/) plugin JACoP (http://rsbweb.nih.gov/ij/plugins/track/jacop.html).

### Immunoelectron microscopy

Human donor retina samples 199-09 and 121-10 were processed for pre-embedding immunolabelling as previously described (88). Ultrathin sections were analyzed with a FEI Tecnai 12 transmission electron microscope. Images were obtained with a charge-coupled device camera (SIS Megaview3; Surface Imaging Systems) acquired by AnalSIS (Soft Imaging System) and processed with Photoshop CS (Adobe Systems).

### HEK293T cell culture

HEK293T cells were cultured in Dulbecco’s modified Eagle’s medium (DMEM), 10% fetal calf serum (FCS, Invitrogen) and 1% penicillin/streptomycin (Invitrogen) at 37°C and 5% CO_2._ Transfections of plasmids were performed with Lipofectamine® LTX and Plus Reagent (Invitrogen), siRNAs were transfected using Lipofectamin RNAiMAX, according to manufacturer’s protocols, respectively.

### Human primary fibroblast cultures

Dermal primary fibroblast lines were expanded from skin biopsies of human subjects (ethics votume: Landesärztekammer Rhineland-Palatinate to KNW). Primary fibroblast lines were mycoplasma negative and cultured in DMEM, 10% FCS and 1% penicillin-streptomycin at 37°C and 5% CO_2_. For ciliogenesis, fibroblasts were starved in OPTIMEM reduced-serum medium (Invitrogen by Thermo Fisher Scientific) for 48 h and processed for immunocytochemistry as previously described (3). The length of Arl13b-positive primary ciliary axonemes were measured directly in images taken with a 63-x objective with a Leica DM6000B deconvolution microscope. For statistical analysis of ciliary length R was applied (https://www.rstudio.com/products/rstudio/download/).

### siRNA knock-down

siRNAs against human USH1C/harmonin (siHarm1 and siHarm2) and non-targeting control siRNA (NTC) were purchased from IDT (TriFECTa^®^ Kit, DsiRNA Duplex, IDT). For knock-downs, HEK293T cells were transfected with harmonin_a1-HA (pBI-CMV4-harm_a1-HA), and 20 nM non-targeted control. 24 h later, cells were lysed in Triton X-100 lysis buffer and protein concentrations were determined using BCA assay. Equal amounts of protein lysates were subjected to SDS-PAGE, followed by Western blotting. Actin was used as a loading control. For quantification, the optical densities of harmonin_a1 (∼ 80 kDa) bands were ascertained and normalized to the appropriate loading control. The percentage (%) of harmonin_a1 expression is shown in relation to harmonin a1-HA transfected cells.

### GST-pull downs

Constructs encoding harmonin domains were cloned in the pDEST17-vector (Gateway cloning system, Invitrogen, USA). The cDNAs encoding *C*-terminal tails of Mm JAM-B tail (amino acids 257-299), Mm β-catenin tail (amino acids 734-782) and filamin A tail (amino acids 2264-2648) were cloned in the vectors pDEST17 and pDEST15 and proteins were bacterially expressed. GST-pull down assays were performed as previously described (15).

### GFP-Trap®

HEK293T cells were transfected with harmonin_a1-HA (pBI-CMV4-harm_a1-HA), GFP-Rhodopsin (pMT3-Rhodopsin-GFP) or GFP (pMT3-Rhodopsin-GFP), respectively, and lysed in Triton X-100 lysis buffer. Protein lysates of transfected cells were precleared using agarose beads. Equal amounts of precleared lysates were made in dilution buffer (10 mM Tris/Cl pH 7.5, 150 mM NaCl, 0.5 mM EDTA). 10% of the diluted amount was used as input. GFP-fused polypeptides were immobilized at BSA blocked-Trap® agarose beads and used for co-precipitation assays according to the manufacturer’s protocol (ChromoTek). Precipitated protein complexes were eluted with SDS-sample buffer at 65°C. Protein complexes were then subjected to SDS-PAGE and Western blot.

### Proximity ligation assay (PLA)

For *in situ* proximity ligation assay the Duolink PLA probes anti-rabbit^PLUS^ and anti-mouse^MINUS^, and the Detection Reagent Red were purchased from Sigma-Aldrich. PLAs were performed as previously described (89, 90) according to manufacturer’s protocol and adapted to our immunohistochemistry protocol for human retinae. As primary antibodies anti-harmonin (H3) and a cocktail of three mAb opsin clones B6-30a1, K16-155, and R2-15) were applied. Negative control was were probed with anti-opsin antibodies and paired with the rabbit- and mouse-IgG-specific oligonucleotide-labelled antibodies. Mounted sections were analyzed on a Leica DM6000B microscope. For quantification, signals from controls were subtracted from signals of antibody combinations in four sections, each.

### Rod outer segment (ROS) enrichment

ROS were purified from 5 pig retinae using a method adapted from (91). Harvested pellets with the ROS were stored at −80°C before use in immunoblot analyses.

### AlphaFold2 modelling

Modelling was done using the hetero-oligomer options of ColabFold (92). Fasta sequence of the respective protein was used from UniProt (UniProt 2020) and a colon between the respective sequences simulated complexes. The following sequences were used: Human USH1C/Harmonin_a1: Gene accession number: Q9Y6N9 harmonin_N-term aa 1-86, harmonin_PDZ1 aa 87-169; harmonin_PDZ2 aa 211-293, harmonin_CC-domain aa 310-377, harmonin_PDZ3 aa 452-537; rhodopsin: Gene accession number: P08100; Adrenergic receptor α-A1: Gene accession number: P35348. Alphafold2 notebook was used in the ColabFold version (https://colab.research.google.com/github/sokrypton/ColabFold/blob/main/AlphaFold2.ipynb#scrollTo=G4yBrceuFbf3) accessed last on March 15, 2022), default options were used but included template mode. Structures were visualized, inspected, and superimposed using PyMOL (Schrödinger, LLC), which was also used to make all figures.

### Statistics

The statistical methods and the significance criteria for gene expression analysis and protein level analysis are listed in corresponding individual legends. Results are shown as means ± SEM of data from at least 3 separate experiments. Two-tailed Student’s t test was used, significance was determined as: *p≤0.05, **p≤0.01.

## Author contributions

KNW, UW conceived, supervised the research project, wrote, reviewed, edited the manuscript. AG, MAN, AS supervised bioinformatics. BF generated RNA-seq libraries. BF, TM, MB, MS, TA, AL analyzed RNA-seq data. AG, LK performed and analyzed bulk RNA-seq. Data collection and analyses were performed by JS (ciliogenesis), MB, TG, KNW (RT-PCR, PLAs, Western blots, immunohistochemistry, TEM), KAW (Western blots), DS, BG (Co-IP, pull-downs) and KF (ROS purification). JF performed Alphafold2 prediction analyses. KS, SK provided human skin biopsies and generated dermal primary fibroblasts. KS did clinical examination of USH1C patients. MMD, US-S, IKK, LAO, MA, JMV, NP provided human donor eyes. Data were discussed with all coauthors.

## Acknowledgments

We wish to thank Ulrike Maas, Elisabeth Sehn, and Gabriele B. Stern-Schneider for their excellent technical assistance. This work was supported by FAUN Foundation (Nuremberg) (UW, KNW), USHER2020 (UW, KNW), the Foundation Fighting Blindness (FFB PPA-0717-0719-RAD; TA-GT-0316-0694-JGU) (UW, KNW), the German Research Council/DFG in the framework of the SPP SPP2127 - Gene and Cell based therapies to counteract neuroretinal degeneration: NA1398/1-1 (KNW), GR4403/5-1 (AG), KO2176/3-1 (SK), WO548/9-1 (UW), European Union Seventh Framework Program under the grant agreements 242013 (TREATRUSH) (UW) and 241955 (SYSCILIA) (UW), ProRetina Foundation Germany (Pro-Re/Seed/Kaplan-Grosche.8-2019) (AG) and NEI-IRP ZIAEY000546 (AS).

## Supplementary material

### Supplemental Figures

**Supplemental Figure S1.**
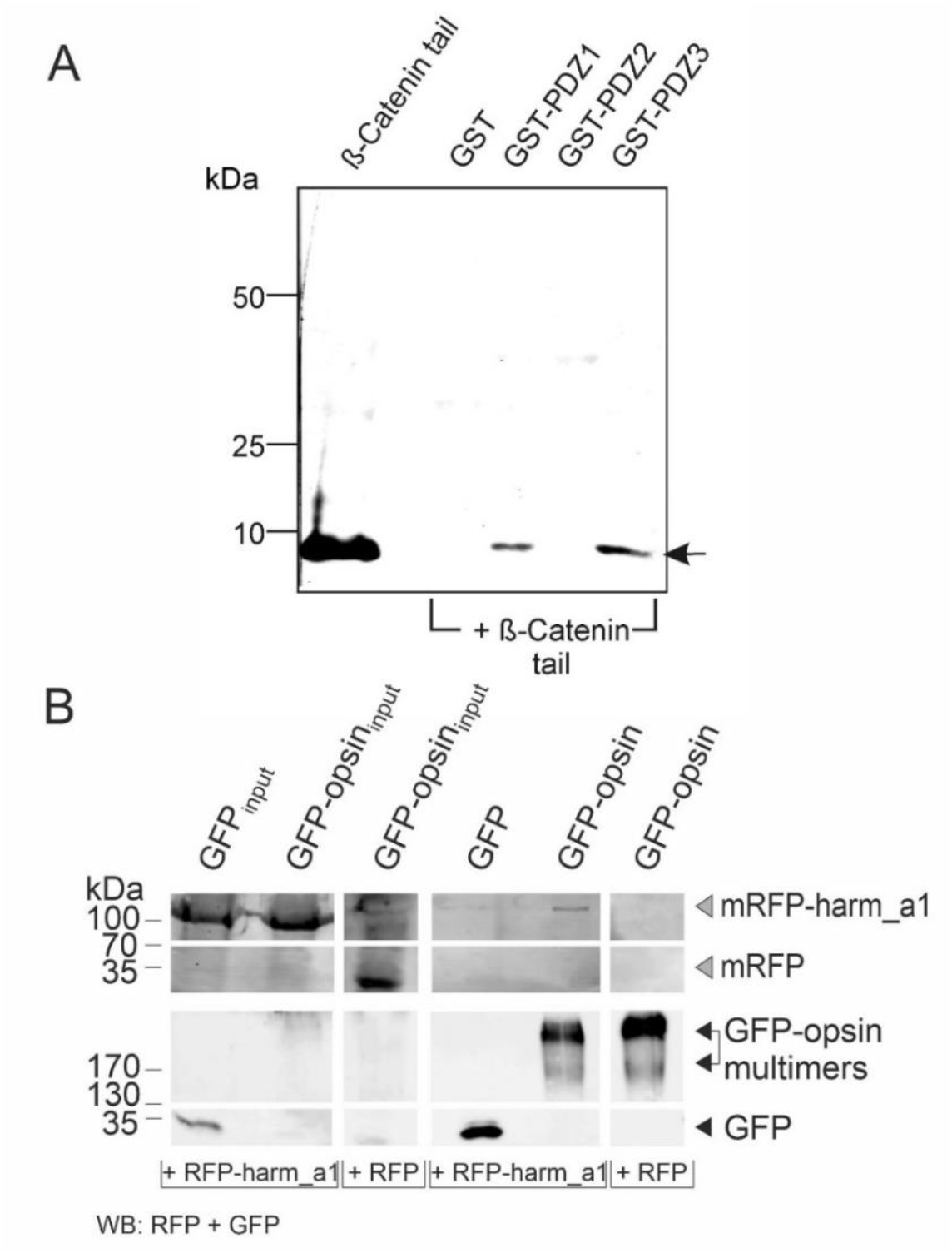
Analysis of novel interaction partners of harmonin. **(A)** GST-pull down of harmonin PDZ domains and β-catenin tail. PDZ1 and PDZ3 of harmonin are interacting with the tail of β-catenin (β-Catenin tail). GST only and harmonin PDZ2 are not interacting with β-catenin. **(B)** GFP-Trap® demonstrate harmonin-RFP interaction with GFP-opsin. RFP-harmonin a1 was precipitated by immobilized GFP-opsin via GFP-beads, but not by GFP alone. RFP only did not interact with GFP-opsin.

**Supplemental Figure S2.**
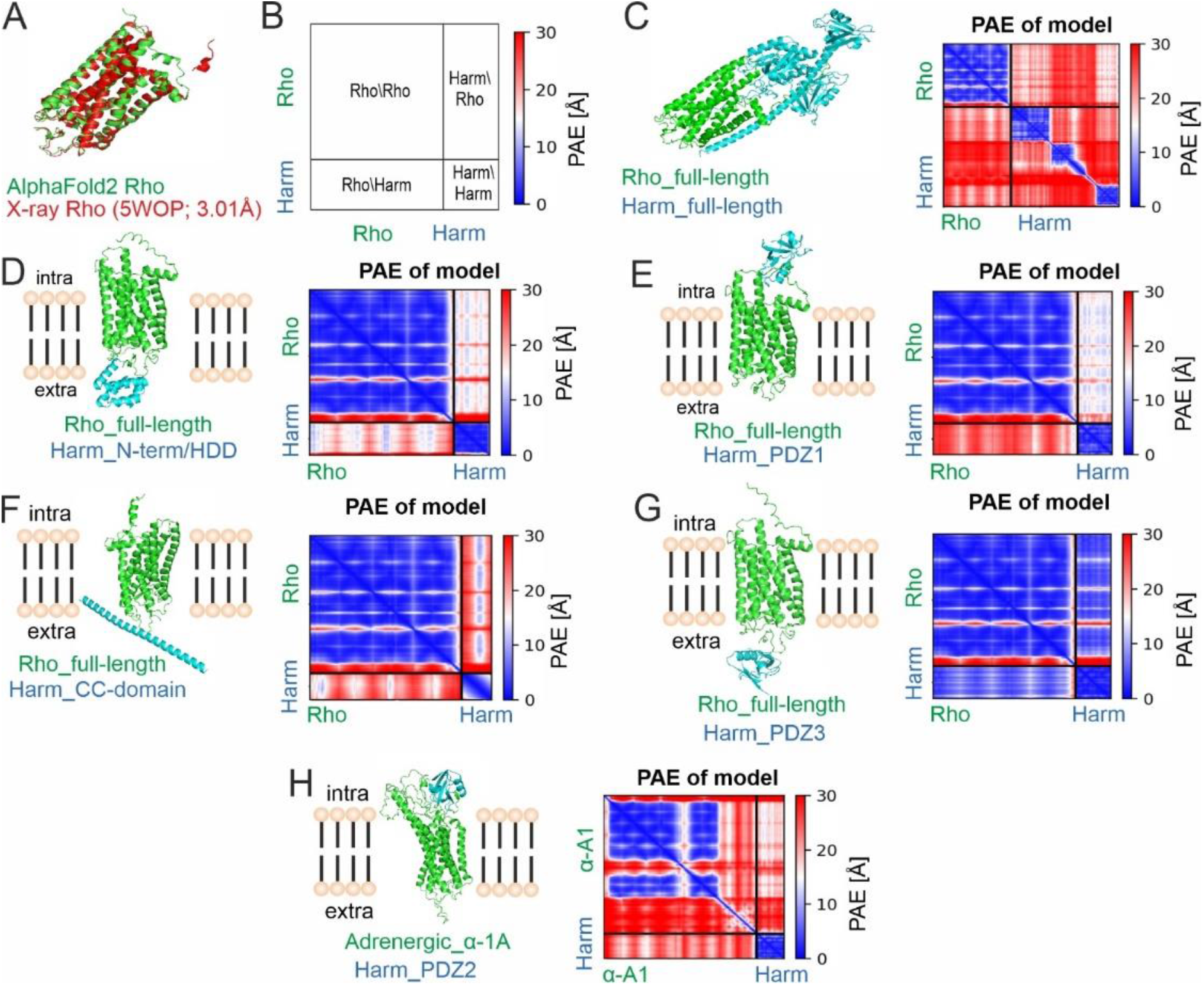
Structure predictions by AlphaFold2. **(A)** The alignment of rhodopsin structures predicted by AlphaFold2 and determined by X-ray crystallography and NMR shows a high degree of concordance, indicating high accuracy of AlphaFold prediction for rhodopsin (Rho) structure based on amino acid sequences. **(B)** Schematic representation of the Predicted Alignment Error (PAE) of a model predicted by AlphaFold. As example: Rho-Harm protein complex. Rectangles lower left and upper right indicate Harm-Rod and Rho-Harm protein complexes, respectively. Heat map illustrates amino acid distances of the two analysed proteins in Å (red: large distance, blue: short distance, predicting protein-protein interaction). (**C**) Structure of the protein complex of full-length rhodopsin (Rho) and full-length harmonin_a1 predicted by AlphaFold2. Predicted Alignment Error (PAE) of the model is high in almost all regions indicating no interaction (red color in heatmap of Ham:Rho and Rho:Harm rectangles). (**D-H**) Structures of the protein complexes of (D) rhodopsin-harmonin_*N-*term/HHD, (**E**) rhodopsin-harmonin_PDZ1, (**F)** rhodopsin-harmonin_CC1, (**G**) rhodopsin-harmonin_PDZ3, and (**H**) full-length adrenergic receptor α-A1 and harmonin_PDZ2 predicted by AlphaFold. AlphaFold predicted low confidence of protein complexes for rhodopsin-harmonin_PDZ1and adrenergic receptor α-A1 and harmonin_PDZ2. For rhodopsin-harmonin_*N-*term/HHD, rhodopsin-harmonin_CC1, and rhodopsin-harmonin_PDZ3 complexes AlphaFold predicted binding of the intracellular protein harmonin to extracellular sites of rhodopsin.

**Supplemental Figure S3.**
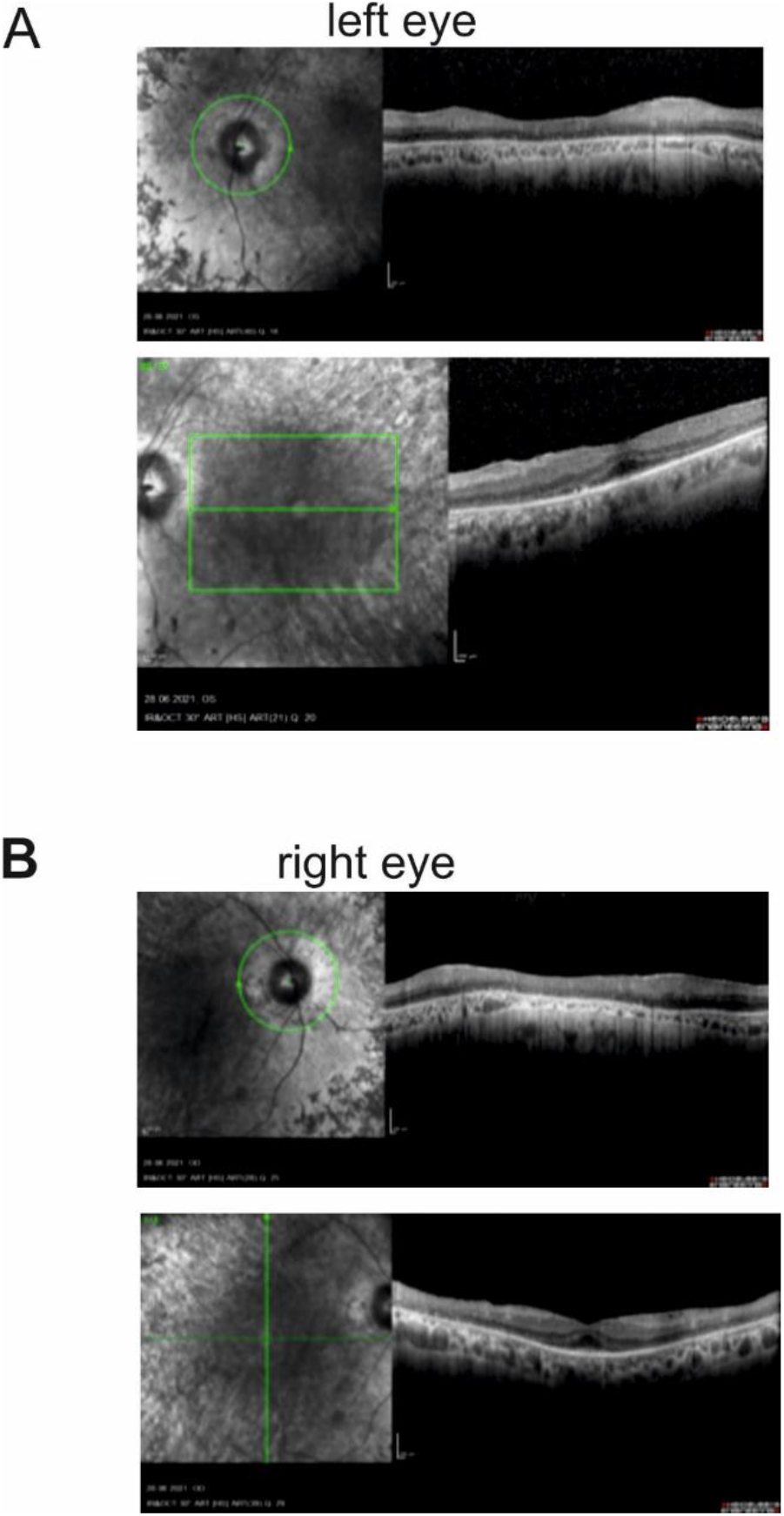
OCT of an USH1C patient. (**A, B**) Optical coherence tomography of the left (A) and right (B) eye of a 47-year old male USH1C patients with confirmed mutations in *USH1C* (c.91C>T;.p.(R31*); c.238dupC;p.(Arg80Profs*69)) showing outer retinal atrophy with photoreceptor degeneration.

### Supplemental Tables

**Supplemental Table S1.**
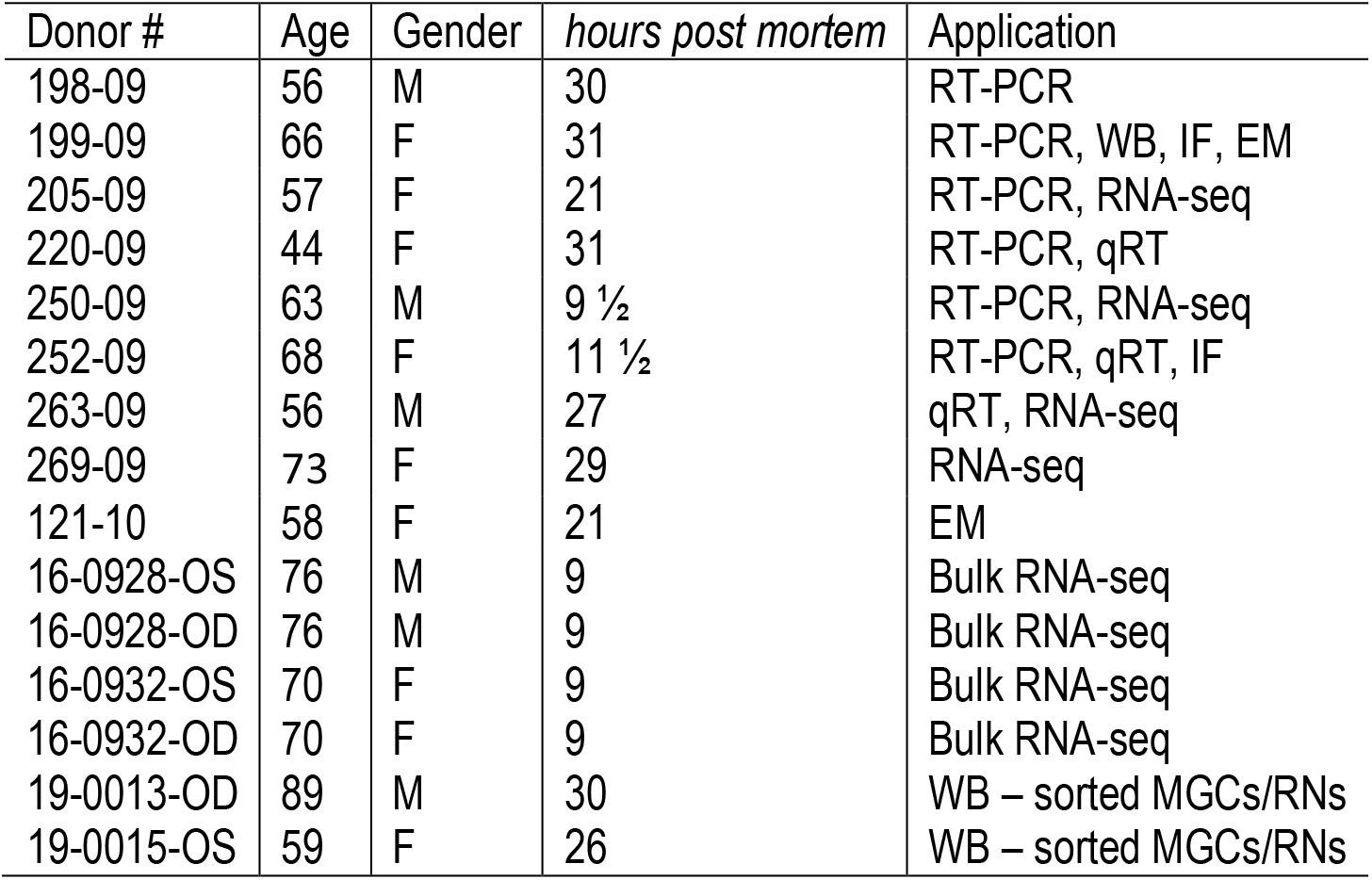
Human retina donors. An internal donor number (#) was assigned to all donors. All donors had no documented history of retinal disease. Abbreviations: M, male, F, female, MGCs, Müller glia cells; RNs, retinal neurons; RT-PCR, reverse transcriptase polymerase chain reaction; qRT, quantitative PCR; WB, Western blot; IF, immunofluorescence microscopy; EM, electron microscopy; age in years.

**Supplemental Table S2. Bulk RNA-seq analyses of *USH1C*/harmonin transcripts in human retina**. List of LSV of *USH1C*/harmonin transcripts in enriched Müller glia cells and retinal neurons (see excel sheets).

**Supplemental Table S3.**
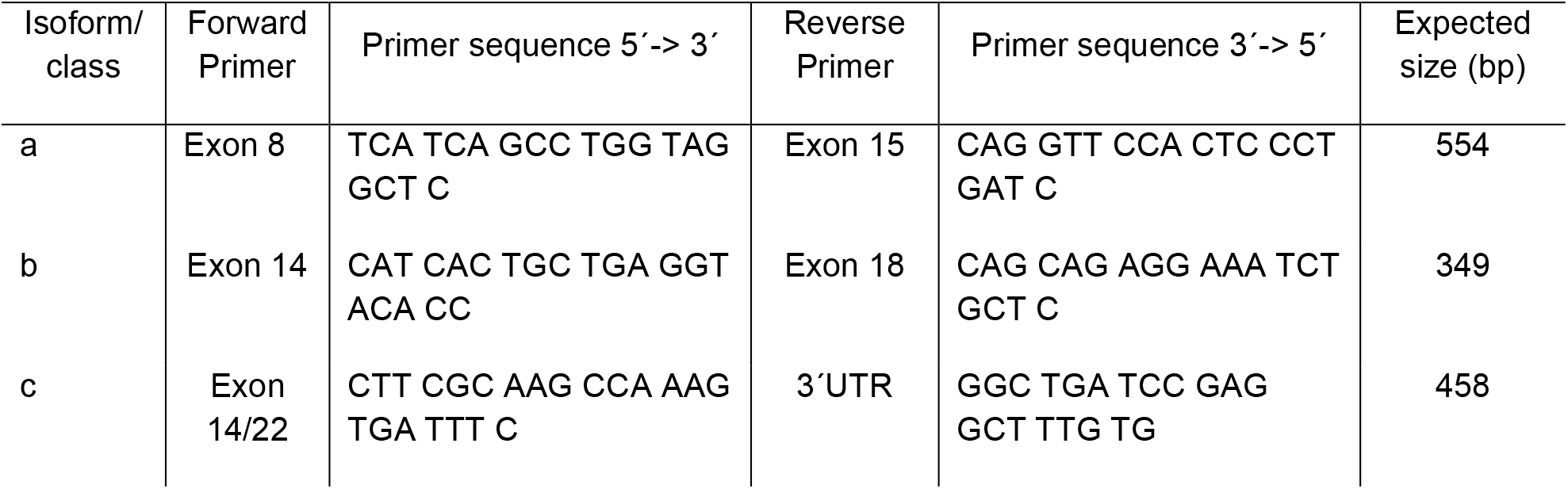
Primers for *USH1C*/harmonin class specific transcripts.

## References

1. Kimberling WJ, et al. Frequency of Usher syndrome in two pediatric populations: Implications for genetic screening of deaf and hard of hearing children. Genet Med 2010. 12:512–516.

2. Friedman TB, et al. Usher syndrome: hearing loss with vision loss. Adv Otorhinolaryngol 2011. 70:56–65.

3. Samanta A, et al. Ataluren for the Treatment of Usher Syndrome 2A Caused by Nonsense Mutations. Int J Mol Sci 2019. 20.

4. May-Simera H, et al. Cilia - The sensory antennae in the eye. Prog Retin Eye Res 2017.

5. Bujakowska KM, et al. Photoreceptor Cilia and Retinal Ciliopathies. Cold Spring Harb Perspect Biol 2017.

6. Fuster-Garcia C, et al. Usher Syndrome: Genetics of a Human Ciliopathy. Int J Mol Sci 2021. 22.

7. Wolfrum U. Protein networks related to the Usher syndrome gain insights in the molecular basis of the disease. In Usher Syndrome: Pathogenesis, Diagnosis and Therapy. S. Ahuja, editor. USA: Nova Science Publishers Inc. 2011. 51–73.

8. Reiners J, et al. Molecular basis of human Usher syndrome: deciphering the meshes of the Usher protein network provides insights into the pathomechanisms of the Usher disease. Exp Eye Res 2006. 83:97–119.

9. Zwaenepoel I, et al. Identification of three novel mutations in the USH1C gene and detection of thirty-one polymorphisms used for haplotype analysis. Hum Mutat 2001. 17:34–41.

10. Verpy E, et al. A defect in harmonin, a PDZ domain-containing protein expressed in the inner ear sensory hair cells, underlies Usher syndrome type 1C. Nat Genet 2000. 26:51–55.

11. Kim E, and Sheng, M. PDZ domain proteins of synapses. Nat Rev Neurosci 2004. 5:771–781.

12. Kremer H, et al. Usher syndrome: molecular links of pathogenesis, proteins and pathways. Hum Mol Genet 2006. 15 Spec No 2:R262-270.

13. Colcombet-Cazenave B, et al. Phylogenetic analysis of Harmonin homology domains. BMC Bioinformatics 2021. 22:190.

14. Adato A, et al. Interactions in the network of Usher syndrome type 1 proteins. Hum Mol Genet 2005. 14:347–356.

15. Reiners J, et al. Scaffold protein harmonin (USH1C) provides molecular links between Usher syndrome type 1 and type 2. Human Molecular Genetics 2005. 14:3933–3943.

16. Yan J, et al. The structure of the harmonin/sans complex reveals an unexpected interaction mode of the two Usher syndrome proteins. Proceedings of the National Academy of Sciences of the United States of America 2010. 107:4040–4045.

17. Boeda B, et al. Myosin VIIa, harmonin and cadherin 23, three Usher I gene products that cooperate to shape the sensory hair cell bundle. EMBO J 2002. 21:6689–6699.

18. Gregory FD, et al. Harmonin inhibits presynaptic Cav1.3 Ca(2)(+) channels in mouse inner hair cells. Nat Neurosci 2011. 14:1109–1111.

19. Gregory FD, et al. Harmonin enhances voltage-dependent facilitation of Cav1.3 channels and synchronous exocytosis in mouse inner hair cells. J Physiol 2013. 591:3253–3269.

20. Yan D, et al. An isoform of GTPase regulator DOCK4 localizes to the stereocilia in the inner ear and binds to harmonin (USH1C). J Mol Biol 2006. 357:755–764.

21. Reiners J, et al. Differential distribution of harmonin isoforms and their possible role in Usher-1 protein complexes in mammalian photoreceptor cells. Invest Ophthalmol Vis Sci 2003. 44:5006–5015.

22. Williams DS, et al. Harmonin in the murine retina and the retinal phenotypes of Ush1c-mutant mice and human USH1C. Investigative Ophthalmology and Visual Science 2009. 50:3881–3889.

23. Sahly I, et al. Localization of Usher 1 proteins to the photoreceptor calyceal processes, which are absent from mice. J Cell Biol 2012. 199:381–399.

24. El-Amraoui A, and Petit, C. Usher I syndrome: unravelling the mechanisms that underlie the cohesion of the growing hair bundle in inner ear sensory cells. Journal of Cell Science 2005. 118:4593–4603.

25. Millan JM, et al. An update on the genetics of usher syndrome. J Ophthalmol 2011. 2011:417217.

26. Phillips JB, et al. Harmonin (Ush1c) is required in zebrafish Muller glial cells for photoreceptor synaptic development and function. Dis Model Mech 2011. 4:786–800.

27. Grotz S, et al. Early disruption of photoreceptor cell architecture and loss of vision in a humanized pig model of usher syndromes. EMBO Mol Med 2022.e14817.

28. Schietroma C, et al. Usher syndrome type 1-associated cadherins shape the photoreceptor outer segment. J Cell Biol 2017. 216:1849–1864.

29. Cowan CS, et al. Cell Types of the Human Retina and Its Organoids at Single-Cell Resolution. Cell 2020. 182:1623–1640 e1634.

30. Xu L, et al. Clarin-1 expression in adult mouse and human retina highlights a role of Muller glia in Usher syndrome. J Pathol 2020. 250:195–204.

31. Bringmann A, et al. Muller cells in the healthy and diseased retina. Prog Retin Eye Res 2006. 25:397–424.

32. Gao H, et al. Muller Glia-Mediated Retinal Regeneration. Mol Neurobiol 2021. 58:2342–2361.

33. Nagel-Wolfrum K, et al. Translational read-through as an alternative approach for ocular gene therapy of retinal dystrophies caused by in-frame nonsense mutations. Vis Neurosci 2014. 31:309–316.

34. Katz Y, et al. Quantitative visualization of alternative exon expression from RNA-seq data. Bioinformatics 2015. 31:2400–2402.

35. Pinelli M, et al. An atlas of gene expression and gene co-regulation in the human retina. Nucleic Acids Res 2016. 44:5773–5784.

36. Scanlan MJ, et al. Isoforms of the human PDZ-73 protein exhibit differential tissue expression. Biochim Biophys Acta 1999. 1445:39–52.

37. Omri S, et al. The outer limiting membrane (OLM) revisited: clinical implications. Clin Ophthalmol 2010. 4:183–195.

38. Golenhofen N, and Drenckhahn, D. The catenin, p120ctn, is a common membrane-associated protein in various epithelial and non-epithelial cells and tissues. Histochem Cell Biol 2000. 114:147–155.

39. Daniele LL, et al. Novel distribution of junctional adhesion molecule-C in the neural retina and retinal pigment epithelium. J Comp Neurol 2007. 505:166–176.

40. Su W, et al. Filamin A is a regulator of blood-testis barrier assembly during postnatal development in the rat testis. Endocrinology 2012. 153:5023–5035.

41. DuChez BJ, et al. Characterization of the interaction between beta-catenin and sorting nexin 27: contribution of the type I PDZ-binding motif to Wnt signaling. Biosci Rep 2019. 39.

42. Ebnet K, et al. The junctional adhesion molecule (JAM) family members JAM-2 and JAM-3 associate with the cell polarity protein PAR-3: a possible role for JAMs in endothelial cell polarity. J Cell Sci 2003. 116:3879–3891.

43. tom Dieck S, and Brandstatter, JH. Ribbon synapses of the retina. Cell Tissue Res 2006. 326:339–346.

44. Blanks JC, and Johnson, LV. Specific binding of peanut lectin to a class of retinal photoreceptor cells. A species comparison. Invest Ophthalmol Vis Sci 1984. 25:546–557.

45. Montell C. Visual transduction in Drosophila. Annu Rev Cell Dev Biol 1999. 15:231–268.

46. Jumper J, et al. Highly accurate protein structure prediction with AlphaFold. Nature 2021. 596:583–589.

47. Evans R, et al. Protein complex prediction with AlphaFold-Multimer. bioRxiv 2021.

48. Chen HY, et al. Primary cilia biogenesis and associated retinal ciliopathies. Semin Cell Dev Biol 2021. 110:70–88.

49. Pan B, et al. Gene therapy restores auditory and vestibular function in a mouse model of Usher syndrome type 1c. Nat Biotechnol 2017. 35:264–272.

50. Aisa-Marin I, et al. The Alter Retina: Alternative Splicing of Retinal Genes in Health and Disease. Int J Mol Sci 2021. 22.

51. Zelinger L, and Swaroop, A. RNA Biology in Retinal Development and Disease. Trends Genet 2018. 34:341–351.

52. Murphy D, et al. Alternative Splicing Shapes the Phenotype of a Mutation in BBS8 To Cause Nonsyndromic Retinitis Pigmentosa. Mol Cell Biol 2015. 35:1860–1870.

53. Ling JP, et al. ASCOT identifies key regulators of neuronal subtype-specific splicing. Nat Commun 2020. 11:137.

54. Sundar J, et al. The Musashi proteins MSI1 and MSI2 are required for photoreceptor morphogenesis and vision in mice. J Biol Chem 2021. 296:100048.

55. Vogel C, and Marcotte, EM. Insights into the regulation of protein abundance from proteomic and transcriptomic analyses. Nat Rev Genet 2012. 13:227–232.

56. Murphy D, et al. The Musashi 1 Controls the Splicing of Photoreceptor-Specific Exons in the Vertebrate Retina. PLoS Genet 2016. 12:e1006256.

57. Yildirim A, et al. SANS (USH1G) regulates pre-mRNA splicing by mediating the intra-nuclear transfer of tri-snRNP complexes. Nucleic Acids Res 2021. 49:5845–5866.

58. Maerker T, et al. A novel Usher protein network at the periciliary reloading point between molecular transport machineries in vertebrate photoreceptor cells. Hum Mol Genet 2008. 17:71–86.

59. Fadl BR, et al. An optimized protocol for retina single-cell RNA sequencing. Mol Vis 2020. 26:705–717.

60. Wu L, et al. Large protein assemblies formed by multivalent interactions between cadherin23 and harmonin suggest a stable anchorage structure at the tip link of stereocilia. J Biol Chem 2012. 287:33460–33471.

61. Grillet N, et al. Harmonin mutations cause mechanotransduction defects in cochlear hair cells. Neuron 2009. 62:375–387.

62. Li J, et al. Mechanistic Basis of Organization of the Harmonin/USH1C-Mediated Brush Border Microvilli Tip-Link Complex. Dev Cell 2016. 36:179–189.

63. Crawley SW, et al. Intestinal brush border assembly driven by protocadherin-based intermicrovillar adhesion. Cell 2014. 157:433–446.

64. Pinette JA, et al. Brush border protocadherin CDHR2 promotes the elongation and maximized packing of microvilli in vivo. Mol Biol Cell 2019. 30:108–118.

65. Testa F, et al. Clinical Presentation and Disease Course of Usher Syndrome Because of Mutations in Myo7a or Ush2a. Retina 2017. 37:1581–1590.

66. Kersten FF, et al. Association of whirlin with Cav1.3 (alpha1D) channels in photoreceptors, defining a novel member of the usher protein network. Investigative Ophthalmology and Visual Science 2010. 51:2338–2346.

67. Hosoi N, et al. Group III metabotropic glutamate receptors and exocytosed protons inhibit L-type calcium currents in cones but not in rods. J Neurosci 2005. 25:4062–4072.

68. Miles A, et al. Usher syndrome type 1-associated gene, pcdh15b, is required for photoreceptor structural integrity in zebrafish. Dis Model Mech 2021. 14.

69. Marshall JD, et al. Alstrom Syndrome. In GeneReviews(R). R.A. Pagon, M.P. Adam, H.H. Ardinger, S.E. Wallace, A. Amemiya, L.J.H. Bean, T.D. Bird, N. Ledbetter, H.C. Mefford, R.J.H. Smith, et al., editors. Seattle (WA): University of Washington, Seattle. GeneReviews is a registered trademark of the University of Washington, Seattle. 1993.

70. Burgoyne T, et al. Rod disc renewal occurs by evagination of the ciliary plasma membrane that makes cadherin-based contacts with the inner segment. Proc Natl Acad Sci U S A 2015. 112:15922–15927.

71. Goldberg AF, et al. Molecular basis for photoreceptor outer segment architecture. Prog Retin Eye Res 2016. 55:52–81.

72. Corral-Serrano JC, et al. PCARE and WASF3 regulate ciliary F-actin assembly that is required for the initiation of photoreceptor outer segment disk formation. Proc Natl Acad Sci U S A 2020. 117:9922–9931.

73. Spencer WJ, et al. Photoreceptor Discs: Built Like Ectosomes. Trends Cell Biol 2020. 30:904–915.

74. Bahloul A, et al. Cadherin-23, myosin VIIa and harmonin, encoded by Usher syndrome type I genes, form a ternary complex and interact with membrane phospholipids. Hum Mol Genet 2010. 19:3557–3565.

75. Bolger AM, et al. Trimmomatic: a flexible trimmer for Illumina sequence data. Bioinformatics 2014. 30:2114–2120.

76. Dobin A, et al. STAR: ultrafast universal RNA-seq aligner. Bioinformatics 2013. 29:15–21.

77. Bray NL, et al. Near-optimal probabilistic RNA-seq quantification. Nat Biotechnol 2016. 34:525–527.

78. Li H, et al. The Sequence Alignment/Map format and SAMtools. Bioinformatics 2009. 25:2078–2079.

79. Robinson JT, et al. Integrative genomics viewer. Nat Biotechnol 2011. 29:24–26.

80. Thorvaldsdottir H, et al. Integrative Genomics Viewer (IGV): high-performance genomics data visualization and exploration. Brief Bioinform 2013. 14:178–192.

81. Grosche A, et al. The Proteome of Native Adult Muller Glial Cells From Murine Retina. Mol Cell Proteomics 2016. 15:462–480.

82. Li B, and Dewey, CN. RSEM: accurate transcript quantification from RNA-Seq data with or without a reference genome. BMC Bioinformatics 2011. 12:323.

83. Schafer N, et al. Complement Components Showed a Time-Dependent Local Expression Pattern in Constant and Acute White Light-Induced Photoreceptor Damage. Front Mol Neurosci 2017. 10:197.

84. Adamus G, et al. Anti-rhodopsin monoclonal antibodies of defined specificity: characterization and application. Vision Res 1991. 31:17–31.

85. Wolfrum U, and Schmitt, A. Rhodopsin transport in the membrane of the connecting cilium of mammalian photoreceptor cells. Cell Motil Cytoskeleton 2000. 46:95–107.

86. Wolfrum U. Centrin-Like and Alpha-Actinin-Like Immunoreactivity in the Ciliary Rootlets of Insect Sensilla. Cell and Tissue Research 1991. 266:231–238.

87. Overlack N, et al. Direct interaction of the Usher syndrome 1G protein SANS and myomegalin in the retina. Biochim Biophys Acta 2011. 1813:1883–1892.

88. Sedmak T, et al. Immunoelectron microscopy of vesicle transport to the primary cilium of photoreceptor cells. Methods Cell Biol 2009. 94:259–272.

89. Bauss K, et al. Phosphorylation of the Usher syndrome 1G protein SANS controls Magi2-mediated endocytosis. Hum Mol Genet 2014. 23:3923–3942.

90. Sorusch N, et al. Characterization of the ternary Usher syndrome SANS/ush2a/whirlin protein complex. Hum Mol Genet 2017.

91. Papermaster DS. Preparation of retinal rod outer segments. Methods Enzymol 1982. 81:48–52.

92. Mirdita M, et al. ColabFold - Making protein folding accessible to all. bioRxiv preprint doi: https://doi.org/10.1101/2021.08.15.456425. 2021

